# CYRI (FAM49) proteins are local inhibitors of Scar/WAVE induced lamellipodia that bind directly to active Rac1

**DOI:** 10.1101/164905

**Authors:** Loic Fort, Jose Batista, Peter Thomason, Heather J. Spence, Jennifer Greaves, Kirsty J. Martin, Kurt I. Anderson, Peter Brown, Sergio Lilla, Matthew P. Neilson, Petra Tafelmeyer, Sara Zanivan, Shehab Ismail, Nicholas C.O. Tomkinson, Luke H. Chamberlain, Robert H. Insall, Laura M. Machesky

**Affiliations:** CRUK Beatson Institute, Switchback Road, Glasgow G61 1BD, UK; University of Glasgow Institute of Cancer Sciences, Switchback Road, Glasgow G61 1BD, UK; Strathclyde Institute of Pharmacy and Biomedical Sciences, University of Strathclyde, 161 Cathedral Street, Glasgow G4 0RE, UK; Hybrigenics Services, 3 Impasse Reille, 75014 Paris, France; WestCHEM, Department of Pure and Applied Chemistry, University of Strathclyde, Glasgow G1 1XL, UK

## Abstract

Actin-based protrusions driving cell migration are reinforced through positive feedback, but it is unclear how the cell restricts the eventual size of a protrusion or limits positive signals to cause splitting or retraction. We have identified an evolutionarily conserved regulator of the protrusion machinery, which we name CYRI (<u>**CY**</u>FIP**-**related <u>**R**</u>ac **i**nteracting) protein. CYRI shows sequence similarity to the Scar/WAVE complex member CYFIP in a <u>D</u>omain of <u>U</u>nknown <u>F</u>unction, DUF1394. CYRI binds specifically to activated Rac1 via a common motif shared with CYFIP, establishing DUF1394 as a new Rac1 binding domain. CYRI-depleted cells have broad, Scar/WAVE-enriched lamellipodia and enhanced Rac1 signaling. Conversely, CYRI overexpression suppresses spreading and dramatically sharpens protrusions into unproductive needles. CYRI proteins use dynamic inhibition of Scar/WAVE induced actin to focus positive protrusion signals and regulate pseudopod complexity. CYRI behaves like a “local inhibitor” predicted and described in widely accepted mathematical models, but not previously identified in living cells.

## Introduction

Cell migration is an ancient and fundamental mechanism by which cells interact with their environment - from seeking nutrients to organizing into specialized tissues. Motile cells have the intrinsic ability to polymerise actin and make protrusions that drive migration. Dozens of proteins regulate the polymerization of actin both spatially and temporally, enabling the cytoskeleton to perform complex and specialised behaviours. The relationship between *de novo* pseudopod generation, pseudopod splitting and retraction during cell translocation is an area of active debate^1^. The 5-protein Scar/WAVE complex is the main driver of Arp2/3 complex induced branched actin networks underlying pseudopod generation. Scar/WAVE is normally autoinhibited until signals from the activated small GTPase Rac1 interacting with the CYFIP subunit (PIR121 in *Dictyostelium*) cause a conformational change leading to interaction with and activation of Arp2/3 complex ^2, 3^. How this activation mechanism is modulated or attenuated is not known, but it is clear that the Scar/WAVE complex is part of a multicomponent network of regulation of which we only currently understand a small fraction of the players. Rac1 is one of the activators of Scar/WAVE, but studies in live cells have revealed much faster dynamics of Scar/WAVE recruitment and turnover at the leading edge than Rac ^4-6^. At least three negative regulators of Arp2/3 complex have been described, Gadkin ^7^ which sequesters Arp2/3, PICK1 (whose role as an Arp2/3 inhibitor is still under debate) ^8, 9^ and Arpin, which mimics the tail of WASP proteins but inhibits rather than stimulating the Arp2/3 complex ^10^. Here, we describe the first negative regulator of Scar/WAVE complex activation by Rac1, CYRI (encoded by the *FAM49* gene), an evolutionarily conserved protein that mimics the Rac1 interaction domain of CYFIP/PIR121 and thus acts as a direct local competitor for activated Rac1 at the plasma membrane.

## Results

### CYRI is an evolutionarily conserved N-myristoylated protein with homology to CYFIPs

We sought new interactors of the Scar/WAVE complex by pulldown of GFP-Nap1 (NCKAP1 in mammals) of the Scar/WAVE complex ^11^ stably expressed in Nap1 knockout *Dictyostelium* cells. Reversible formaldehyde crosslinking in cellulo^12^ stabilised transient interactions and subsequent GFP-trap immunocapture recovered Scar/WAVE complex members Scar/WAVE, ABI, HSPC300 and PIR121 (CYFIP1/2 in mammals). Among other interactors, we noticed an associated uncharacterized protein with homology to human FAM49 (for <u>fam</u>ily of unknown function <u>49</u>) (Fig 1A and Table S1). Even though subsequent attempts to immunoprecipitate FAM49 with the Scar/WAVE complex in the absence of crosslinking showed that this interaction likely indirect, FAM49 caught our attention for two reasons. Firstly, it is highly conserved across evolution, hinting at a fundamental function (Supplementary Fig.1A). *FAM49* is present in all of the major eukaryotic superfamilies (as defined by Keeling *et al*.^13^)-unikonts, chromalveolates, excavates and at least one example in plants and is roughly co-conserved with the Scar/WAVE complex (Supplementary Fig.1B). Secondly, both Pfam and InterPro identified FAM49 as sharing a DUF1394 domain with CYFIPs (Fig. 1B-D and Supplementary Fig.1C) suggesting a functional linkeage with the Scar/WAVE complex. FAM49 is basically a DUF1394, while CYFIPs have a cytoplasmic fragile X interaction domain, previously implicated in neuronal dendritic spine function ^14^ (Fig. 1B). We renamed FAM49 to a more functional name, CYRI for <u>**CY**</u>FIP-related <u>**R**</u>ac1 **i**nteracting <u>**protein**</u>, due to the functions described below. In mammals, the two isoforms are CYRI-A (FAM49-A) and CYRI-B (FAM49-B). We henceforth refer to the FAM49 protein and the human gene as CYRI.

**Figure 1:**
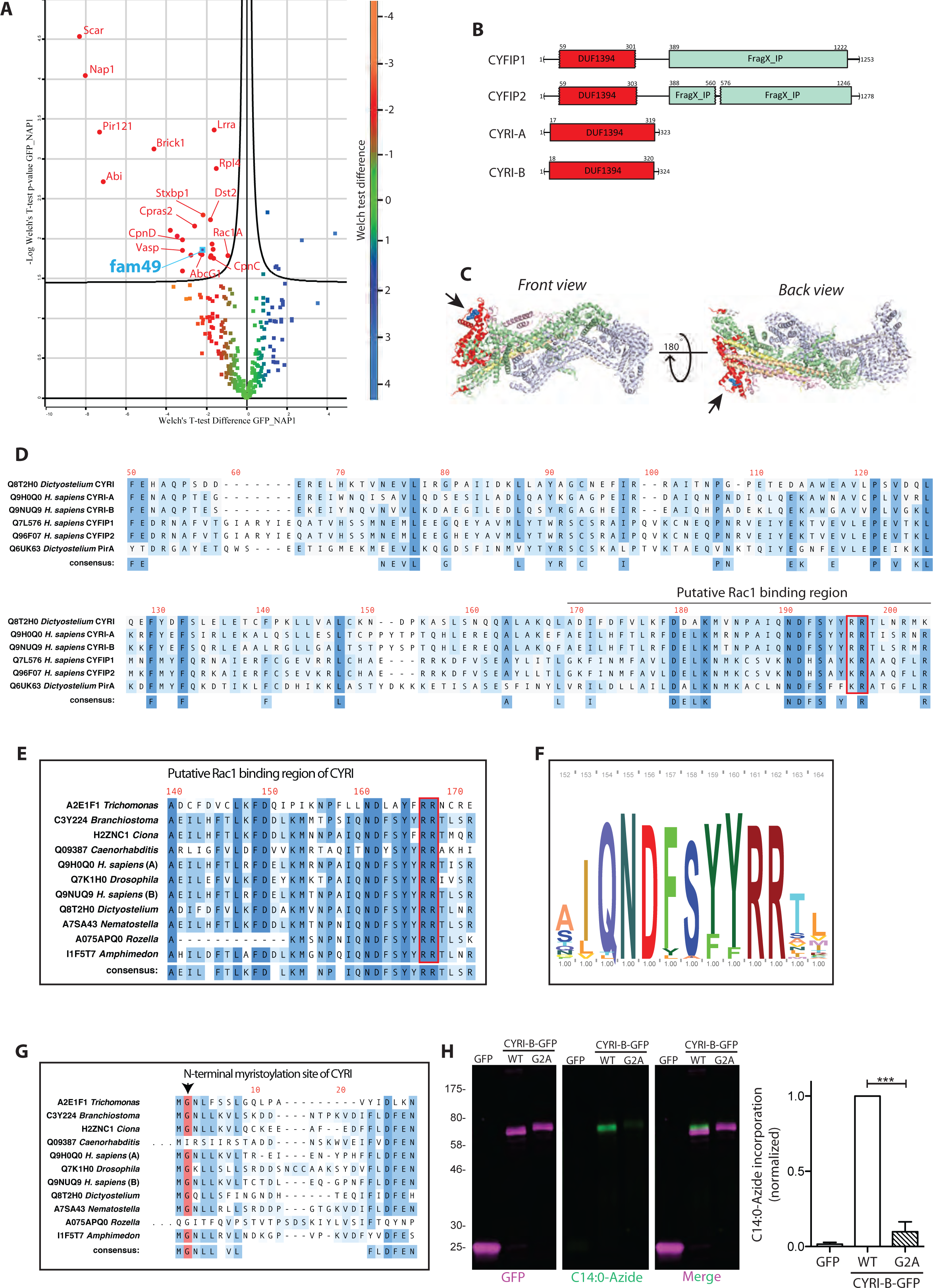
CYRI-B is a highly conserved protein related to CYFIP/ PIR121 and defined by a DUF1394. A – Volcano plot illustrating pooled results from four LC-MS/MS experiments. *D. discoideum* Ax3 wild-type or *napA* knockout *Dictyostelium* cells were stably transfected with GFP or GFP-NAP1 respectively. Following crosslinking using formaldehyde, GFP-tagged proteins were immunoprecipitated and run on a Coomassie gel for LC-MS/MS analysis. Color-coding was done on the basis of the Welch test difference. Curved line represents the 5% false discovery rate threshold (Welch’s test) and enriched proteins are represented with red dots. Identified interactors are labeled with gene symbols and presented in **Table 1** with additional information. B-Comparison between human CYFIP1/2 and CYRI-A/B organization showing amino acid numbers and domains. Common DUF1394 domain (Pfam PF07159) is red whereas CYFIP1/2 C-terminal cytoplasmic Fragile X Mental Retardation FMR1-interacting domain (FragX-IP, Pfam PF05994) is light green. C– Two views of ribbon crystal structure of the Scar/WAVE complex (Chen et al. PDB 3P8C). NCKAP1 in lilac, CYFIP1 in light green and red, Scar/WAVE in peach, HSPC300 in yellow and ABI1 in orange. DUF1394 is red, with putative Rac1 interaction residues blue and highlighted by arrows. D-Sequence alignment of the DUF1394 of CYFIP1/2/Pir121 (PirA) and CYRI-A/B. CYFIP1 Arg190 (position 197 from the alignment consensus) is important for CYFIP1 mediated Rac1-binding. Uniprot accession numbers are listed. E– Sequence alignment of the putative Rac binding domain of CYRI in different organisms. CYFIP Lys189 and Arg190 are well conserved in CYRI (Arg160 and Arg161) and are highlighted in red. F – HMM consensus Pfam alignment of the DUF1394 near the conserved arginine. Letter height divides stack height according to the letter frequency across a seed dataset composed of 38 sequences. G – Sequence alignment covering the N-terminal region of CYRI from representative evolutionarily diverse eukaryotes. The glycine in the 2^nd^ position (highlighted red) is a putative myristoylation site. H– CLICK chemistry analysis of the glycine 2 of CYRI-B. Myristoylation event was labeled in HEK293T cells and measured by incorporation of myristate-azide (green) in GFP-wt (magenta) or G2A CYRI-B mutant (magenta), following GFP immunoprecipitation. Quantification of 3 independent experiments is shown in H. Bar plot shows mean and SEM. One way ANOVA with Tukey test was applied and *** <0.001. Color code represents the number of entries with identical amino acids at each position for D, E, G.

The DUF1394 domain of CYFIPs partly overlaps with the Rac1 interaction site on Scar/WAVE Complex, in particular Arg190 in CYFIP1, which was demonstrated to be important for Rac1 binding^2^. The analogous Arg 161 of CYRI (Fig. 1C **dark blue balls, D red box)** is part of a highly conserved 33-amino acid stretch showing sequence similarity >75% across diverse phyla (Fig. 1D-F).

Previous unbiased mass spectrometry analyses had identified lipid modification of CYRI ^15-17^. The N-termini of CYRI show conservation of a putative myristoylation site at glycine 2 (Fig. 1G), which is not conserved in CYFIPs. We confirmed the myristoylation of CYRI-B by assessing the incorporation of myristate analogue (C14:0-azide) onto the glycine using CLICK chemistry *in cellulo*. Site-directed mutagenesis of this glycine to alanine totally abolishes the CLICK signal (Fig. 1H). The myristoylation at position 2 implies a role for CYRI at the plasma membrane^18^, where Rac1 and the Scar/WAVE complex act to catalyse lamellipodial growth.

### CYRI interacts directly with activated Rac1 and negatively regulates Rac1-Scar/WAVE Complex

The provocative homology of CYRI with CYFIP, in a region that is key for interaction with active Rac1, led us to investigate binding between CYRI-B and active Rac1. A blind yeast two-hybrid screen using active Rac1^G12V^ as bait retrieved CYRI-B from various mouse and human cDNA libraries (Fig. 2A and Supplementary Fig. 2A). The core interacting sequence of CYRI-B was mapped to a central domain encompassing amino acids 30-236 (hereafter referred to as Rac Binding Domain RBD) and tested for Rac1 interaction. GFP-RBD expressed in CHL1 human melanoma cells interacted selectively with purified recombinant active GST-Rac1^Q61L^ but not with GST-Rac1^WT^. Mutation of either Arg160 or Arg161 of GFP-RBD to an acidic amino acid abrogated this interaction (Fig. 2B-C). Thus CYRI-B RBD interacts with active Rac1 and in common with CYFIP, two conserved basic residues in the DUF1394 are important for this interaction. This implies a signal-regulated interaction between active Rac1 and CYRI, in the same way that Rac1 interacts with Scar/WAVE complex.

**Figure 2:**
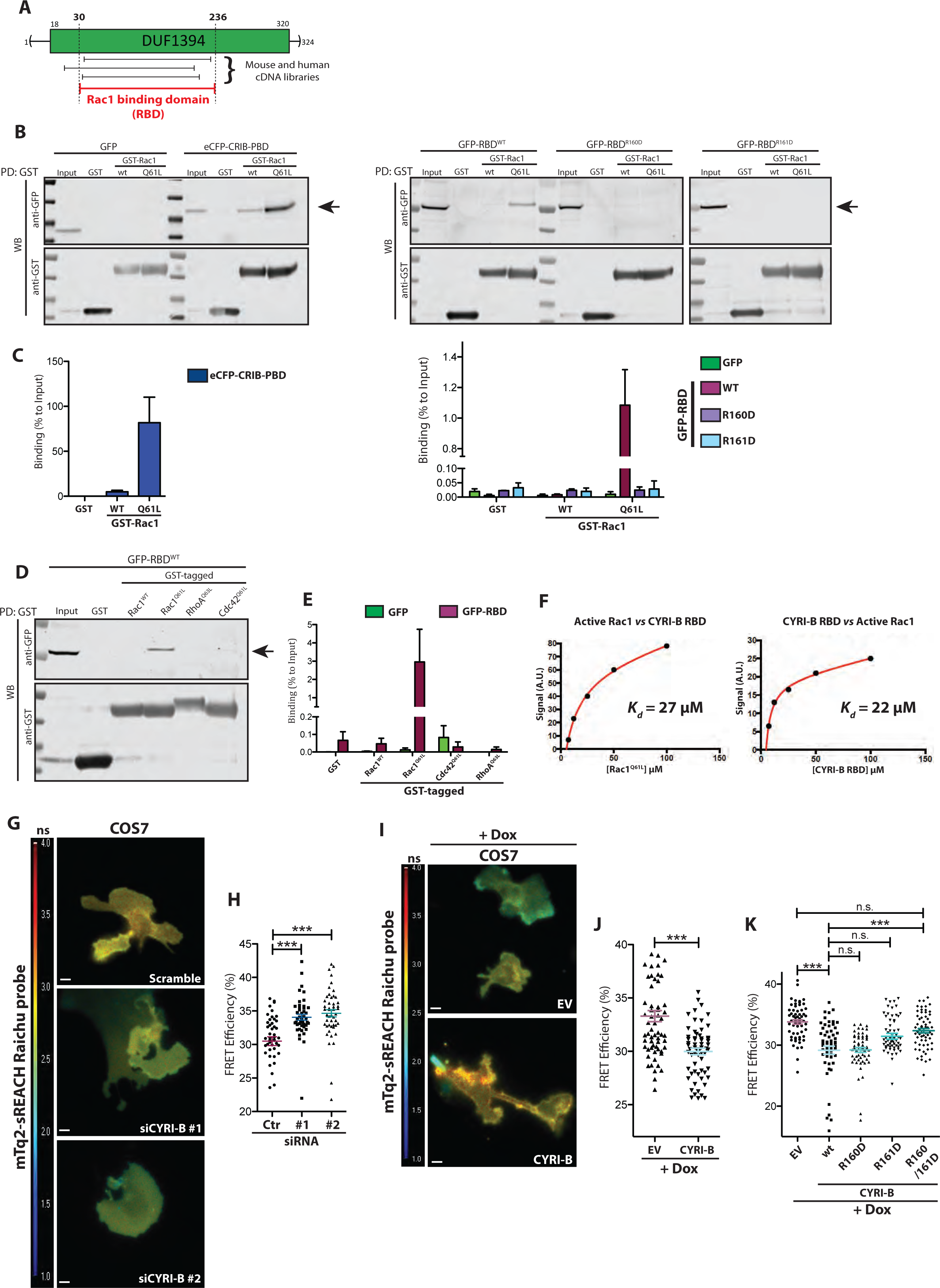
CYRI-B interacts with active Rac1 via its DUF1394 domain. A - Schematic representation of clones obtained by yeast two-hydrid screening various mouse and human cDNA libraries against active Rac1^G12V^. High-confidence interacting regions were used to define a core Rac1-binding domain (RBD) of CYRI-B. (Details in Supplementary Fig. 2A). B - Western blot images from pulldown of GST control, GST-Rac1^WT^ or GST-Rac1^Q61L^ immobilized on beads, mixed with lysate from CHL1 cells expressing either GFP alone, eCFP-CRIB-PBD, GFP-RBD^WT^, GFP-RBD^R160D^ or GFP-RBD^R161D^. Binding (arrow) was quantified relative to the input using densitometry (C). D - Pulldown of GST control, or GST-tagged Rac1^WT^, Rac1^Q61L^, RhoA^Q63L^, Cdc42^Q61L^ immobilized on beads, mixed with lysate from CHL1 cells expressing GFP alone or GFP-RBD^WT^. Binding (arrow) was quantified relative to the input using densitometry (E). F – SPR binding curves of the interaction between purified recombinant Rac1^Q61L^ and CYRI-B-RBD. Left: GST-CYRI-B was immobilized on anti-GST surface vs increasing concentrations of Rac1^Q61L^. Right: His-Rac1 was immobilized on NTA surface vs increasing concentrations of CYRI-B RBD. Analyte injections were performed at five different concentrations and steady state analyses used Biacore evaluation software 3.0, fitting to a simple 1:1 binding model. G-K) FLIM imaging and quantification of CYRI-B knockdown (G-H) or CYRI-B-overexpression after doxycycline induction (I-J) in COS-7 cells. The jet2 color code shows the average lifetime of the mTq2-sREACH Raichu probe, spanning 1-4 ns (blue to red). K) FRET efficiency of COS7 cells overexpressing wt, R160D, R161D or double mutant R160/161D CYRI-B after doxycycline induction. H) *(Ctr n=46, siRNA#1 n=46, siRNA#2 n=48). One-way ANOVA with Dunn’s multiple comparison test was performed. n.s. = not significant p>0.05; *** p≤0.001* J) *(Dox_Ctr n=62, Dox_CYRI-B n=62). Mann Whitney test was performed. K)(EV=59, wt=62, R160D=57, R161D=61, R160/161D=63). One-way ANOVA with Dunn’s multiple comparison test was performed. n.s. p>0.05; *** p≤0.001* All data shown in this figure are representative of at least 3 independent experiments unless specified. Scale bar = 10μm Bar and scatter plots show data points with mean and SEM.

DUF1394 sequences define a new Rac1 binding sequence, as they show no homology to CRIB (<u>C</u>dc42 <u>R</u>ac interaction <u>b</u>inding) motifs commonly found in GTPase binding partners ^19^ or, to our knowledge, any previously described Rac1 interaction sequence. Scar/WAVE complex is specifically regulated by Rac proteins, whereas CRIB motifs generally interact equally with Rac and Cdc42 ^19^. We therefore used GST-pulldowns with purified recombinant active RhoA, Rac1 and Cdc42 to probe the specificity of the interaction of CYRI with Rac1. CYRI-B RBD showed clear interaction with Rac1^Q61L^ but did not interact with constitutively active RhoA^Q63L^ or Cdc42^Q61L^ (Fig. 2D-E and Supplementary Fig. 2B). We tested the Rac1 CYRI-RBD interaction directly using purified recombinant proteins and surface plasmon resonance. Immobilised Rac1 ^Q61L^ specifically bound to purified CYRI-B RBD with a *K_d_* of 27 μM and the reverse assay, with CYRI-B RBD immobilised returned a similar *K_d_* of 22 μM (Fig. 2F). Thus, we conclude that CYRI-B RBD directly interacts with purified active Rac1 and shows specificity for Rac1 over Cdc42 and RhoA. The relatively low affinity of CYRI-B for Rac1 compared with PAK-CRIB (K*^d^* = 0.05 μM ^20^) is consistent with the low affinity of Rac1 for Scar/WAVE complex (1-10 μM ^2^) and the need for a dynamic interaction at the cell leading edge. Furthermore, Scar/WAVE complexes act in clusters at the plasma membrane ^21, 22^, where CYRI likely modulates coincidence detection of activation signals^21, 22^. We thus conclude that CYRI-B selectively binds to activated Rac1, similarly to CYFIP1^2^, thus defining the DUF1394 as a new dynamic detector of activated Rac1 and potential modulator of the coincidence detection by Scar/WAVE complex.

To test our hypothesis that CYRI-B would modulate the Rac1 and Scar/WAVE dynamic interaction in cells, we used a dark acceptor Raichu FRET probe ^23, 24^ to measure activity of Rac1 in cells where we perturbed CYRI-B. Depleting CYRI-B by CrispR-Cas9 mediated knockout in CHL1 melanoma cells or by siRNA in COS7 cells (Supplementary Fig. 2C-D) caused similar increases in the FRET efficiency of the mTq2-sREACH Raichu probe (Fig. 2G-H and Supplementary Fig. 2E-F). This indicates an increase in Rac1 signaling in the absence of CYRI-B. Conversely, inducible overexpression of CYRI-B drove a decrease in the Rac1 activity signal of the probe (Fig. 2I-J and Supplementary Fig. 2G-H). This drop in Rac1 activation signal upon CYRI-B expression was fully reversed by mutation of Arg160 and Arg161 to aspartic acid **(R160/161D mutant**, Fig. 2K). Thus, as well as interacting with Rac1 directly, CYRI-B buffers Rac1 activity at the plasma membrane. This suggests that CYRI-B is a negative regulator of Rac1 and Scar/WAVE induced actin assembly in cells, by binding to Rac1 and lowering its activity, in competition for the interaction between CYFIP and active Rac1.

### CYRI acts locally in pseudopods to restrict lamellipodia protrusions

Since CYRI-B is a negative regulator of Rac1 activity at the plasma membrane, depletion or overexpression should oppose lamellipodia protrusions. Indeed, while *CYRI-B* knockout cells showed a normal growth rate in culture (Supplementary Fig. 3A), they showed severe cytoskeletal alterations and frequently protruded unusually large and broad lamellipodia highly enriched in WAVE2 (Fig. 3A-B), cortactin and the Arp2/3 complex (Supplementary Fig.3B-E) and an increased area of spreading (Fig. 3C). Similar phenotypes were also observed using CYRI-B siRNA-treated COS-7 cells (Fig. 3 D-G and Supplementary Fig. 2D). CYRI-B knockdown cells also have a more circular perimeter than their control counterparts (Fig. 3G), mimicking the classical fried-egg shape observed following Rac1 hyperactivation ^25^. Expression levels of CYFIP1, NCKAP1 and WAVE2 are not detectably altered in *CYRI-B* knockout cells (Supplementary Fig. 3F), suggesting that loss of CYRI-B does not drive lamellipodia by increasing overall levels or stability of the Scar/WAVE complex.

**Figure 3:**
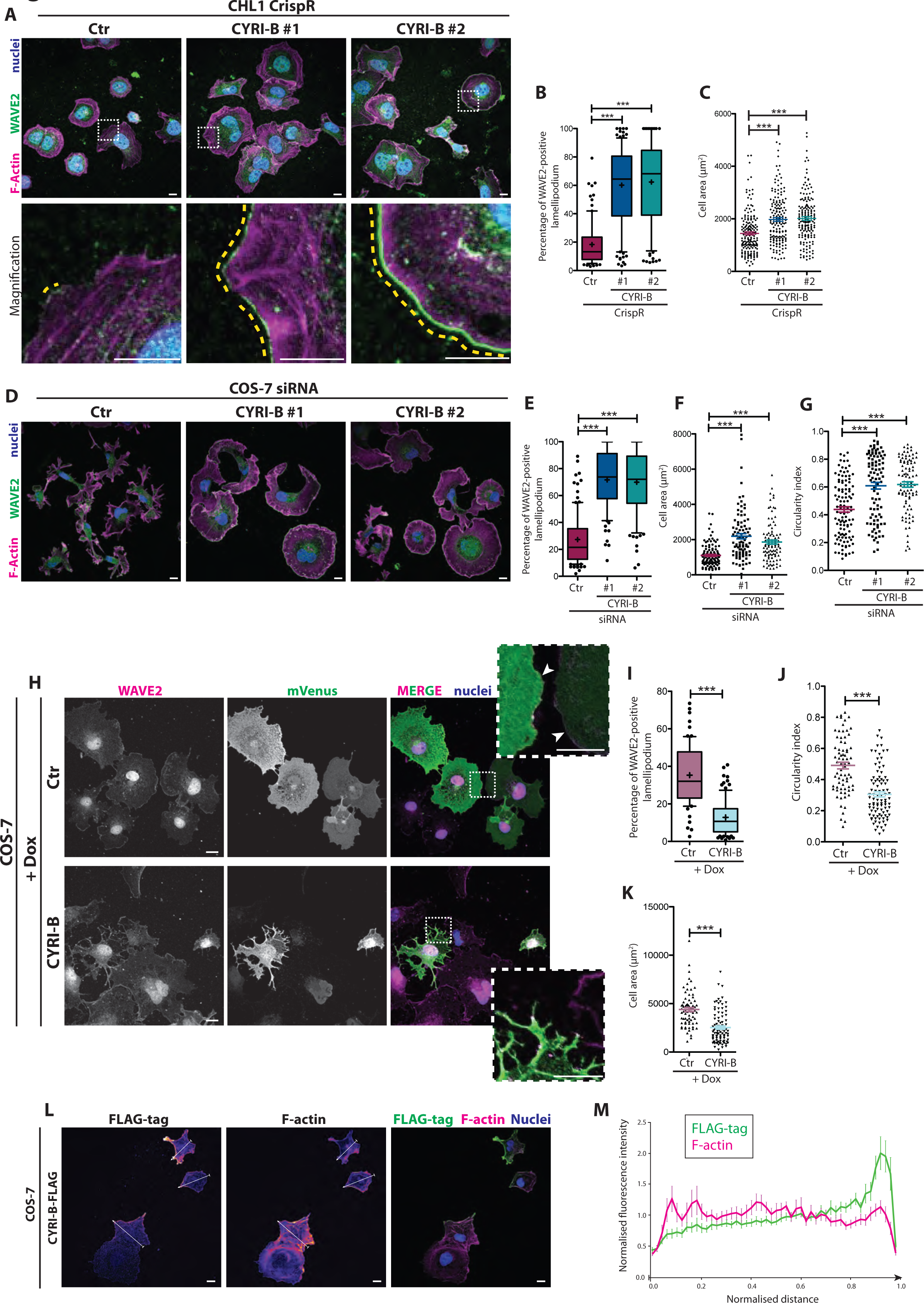
CYRI-B opposes lamellipodial protrusion. A-C) Immunofluorescence (A) of control (Ctr) or each of two independent *CYRI-b* CrispR knockout CHL1 cells plated on collagen and stained for WAVE2 (green), DNA (blue) and F-actin (magenta). Box insets show a magnified field below each panel. The ratio of extension of WAVE2 staining (yellow dotted line) *vs* the total cell perimeter shown in B is a read-out of the extent of the cell edge devoted to lamellipodium. Quantification of cell area after 4h of spreading on collagen is shown in C. *(Ctr n=89, CrispR#1 n=109, CrispR#2 n=97) One-way ANOVA with Dunn’s multiple comparison test. *** p≤0.001.* D-G) Immunofluorescence of control or *CYRI-b* knockdown COS-7 cells plated on laminin for 1h and stained for WAVE2 (green) DNA (blue) and F-actin (magenta) (D). WAVE2 ratio is plotted in E. Cell area and circularity index were measured based on the phalloidin staining and plotted in F and G respectively. *(Ctr n=111, siRNA#1 n=95, siRNA#2 n=96). One-way ANOVA with Dunn’s multiple comparison test. *** p≤0.001.* H-K) Immunofluorescence of doxycycline-induced control or CYRI-B overexpression in COS-7 cells plated on fibronectin for 4h and stained for WAVE2 (magenta), DNA (blue) and GFP (green). Inset panels are a magnified view of the white dashed field. WAVE2 ratio and shape parameters were measured and reported in I and J respectively. Cell area was measured based on the phalloidin staining and plotted in K *(Dox_Ctr n=73, Dox_CYRI-B n=93). Mann Whitney test. *** p≤0.001.* L-M) Immunofluorescence of COS-7 cells transfected with CYRI-B-FLAG and plated on fibronectin-coated coverslips, fixed after 4h and stained for FLAG-tag (green) and F-actin (magenta). White lines show example cross-sections quantified in (M). (M) Quantified normalized fluorescence intensity plot running across 17 representative cells and ending at the protrusive end (normalized distance: 1= protrusive end and 0=opposite end). All data shown in this figure are representative of at least 3 independent experiments. Scale bars = 10μm Bar plots show data points with mean and SEM. Whisker plots show 10-90 percentile and mean (cross).

To determine whether Rac1 was required for the exaggerated lamellipodial phenotype of *CYRI-B* knockout cells, we co-depleted Rac1 and CYRI-B from fibroblasts cultured from a conditional inducible Rac1 knockout mouse ^26^. We used ROSA26-Cre::ER^T2+^;p16Ink4a^−/−^, Rac1^fl/fl^ mouse tail fibroblasts, treated or not with hydroxytamoxifen (to induce deletion of Rac1) and then with siRNA against CYRI-B. Deletion of Rac1 led to a spindle-shaped morphology and a loss of lamellipodia as previously described ^27-29^. Loss of CYRI-B did not cause excessive lamellipodia or altered circularity in Rac-deleted cells (Supplementary Fig. 3G-J). Thus, Rac1 is absolutely required for CYRI-B driven actin reorganisation. Overall, therefore, *CYRI-B* knockout cells show excessive Scar/WAVE driven lamellipodia, caused by Rac1 activation, leading to an increase in spread area. This supports a role for CYRI-B as a buffer of Rac1 activity at the leading edges of cells, regulating activation of the Scar/WAVE complex.

If CYRI is a negative regulator of Rac1 and Scar/WAVE, then increasing expression should oppose lamellipodia. Most cell lines did not tolerate CYRI-B overrexpression, but a doxycycline inducible system in COS-7 cells with a co-expressed membrane-targeted mVenus provided up to 5-fold CYRI-B enrichment after 48h of induction (Supplementary Fig. 2G). Overexpressors frequently appeared smaller, lacked lamellipodia and were either not spread at all or (around 50%) showed highly complex fractal pseudopods with loss of lamellipodia, and consequently decreased circularity. WAVE2 staining at the cell edge was also greatly diminished (Fig. 3H-K and Supplementary Fig. 3K-N). We therefore conclude that overexpression of CYRI-B opposes lamellipodia assembly, maintenance and spreading, all of which require constant polymerization of actin to support retrograde flow. The phenotype is however, different from loss of Rac1 (e.g. see Supplementary Fig. 3H), which makes cells become spindle-shaped^27-29^. This suggests that CYRI-B has a more specific role than Rac1, involving negative regulation of the Rac1 - Scar/WAVE complex pathway at the plasma membrane.

We next localised CYRI-B within cells. No commercially available antibodies gave a reliable signal (not shown), and GFP-CYRI-B or CYRI-B-GFP was not tolerated by cells, nor showed any specific localization. However, low levels of CYRI-B tagged with FLAG, which is much smaller than GFP (Supplementary Figure 3O) revealed a striking peripheral localization of CYRI-B similar to WAVE2, and a gradient of CYRI-B localization across the cell (Fig. 3L-M). CYRI-B levels were at their peak in areas where lamellipodia protrusions were visible by actin staining. This localization suggests that CYRI-B is recruited by the same signals as the Scar/WAVE complex to areas of active protrusion.

### CYRI focuses actin assembly in leading pseudopods to promote plasticity of migration

As lamellipodia and Rac1 – Scar/WAVE are major regulators of cell migration, we explored a role for CYRI-B in 2D migration and 3D invasion. CHL1 human melanoma cells were sparsely seeded onto fibronectin-coated dishes and viewed for 17h by timelapse microscopy. *CYRI-B* knockout cells migrated on average 1.5 fold faster and travelled a greater aggregate distance (Fig. 4A-B). Furthermore, CYRI-B knockouts were more invasive and migrated further in two different Matrigel invasion assays (Supplementary Fig. 4A-D). Thus CYRI-B controls migration and may restrict cancer invasion, via its regulation of Rac1 and Scar/WAVE complex^30^.

**Figure 4:**
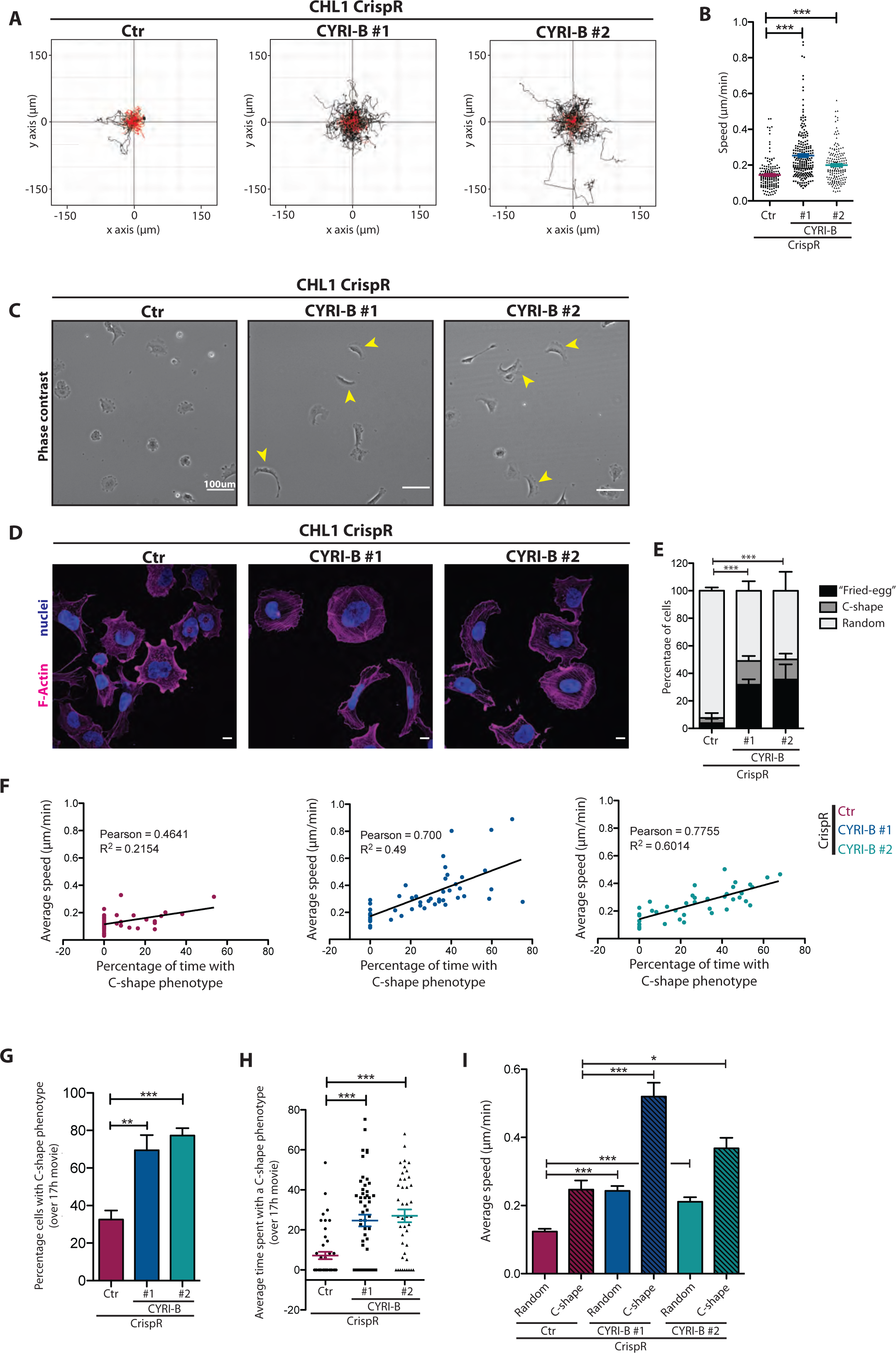
Regulation of membrane protrusion by CYRI-B affects cell migration. A) Spider plots for random migration tracks of control (Ctr) or CYRI-B CrispR knockout CHL1 cells plated on fibronectin. Black tracks represent cells with a travelled distance greater than 100 μm during 17h and red less than 100μm. Average speed is plotted in B. *(Ctr: n=161, CrispR#1 n=228, CrispR#2 n=178). One-way ANOVA with Dunn’s multiple comparison test. *** p≤0.001.* C) Still phase contrast pictures from a random migration assay of control (Ctr) or *CYRI-B* CrispR knockout CHL1 cells plated on fibronectin. Yellow arrowheads denote C-shaped cells. Scale bar = 100μm (See also Supplementary Movie 1) D) Immunofluorescence pictures of control (Ctr) or *CYRI-B* CrispR knockout CHL1 cells plated on collagen, and stained for F-actin (magenta) and DNA (blue). Cells were classified into 3 categories according to their morphology (Fried-egg, C-shape, Random) and the percentage of each population is shown in E. *(Ctr n=276, CrispR#1 n=216, CrispR#2 n=210). Two-tailed Chi-square test (95% confidence). *** p≤0.001.* Scale bar = 10μm F) Correlation between cell shape and average speed in control (Ctr) or *CYRI-B* CrispR knockout CHL1 cells. Pearson coefficient and R^2^ value are shown for each condition. *(Ctr: n=45, CrispR#1 n=53 CrispR#2 n=42).* G-I) Analysis of random migration assay movies to show percentage of cells presenting a C-shape and time spent with this phenotype. G - Cochran-Mantel-Haenszel test **<0.001; ***<0.0005 H - *One-way ANOVA with Dunn’s multiple comparison test was performed. *** p≤0.001.* *(Ctr: n=45, CrispR#1 n=53 CrispR#2 n=42).* I) Speed of control (Ctr) or *CYRI-B* CrispR knockout cells displaying a random or a C-shape. *One-way ANOVA with Dunn’s multiple comparison test was performed. * p≤0.05; *** p≤0.001* All data shown in this figure are representative of at least 3 independent experiments. Bar and scatter plots show data points with mean and SEM

To analyse the mechanisms of CYRI-B migration suppression, we studied the shapes of the faster moving cells in 2D random migration assays. Time-lapse microscopy revealed that *CYRI-B* knockout cells frequently adopted a C-shape when migrating across a flat 2D surface, with a broad spread lamellipodium spanning approximately the front half of the cell perimeter and a convex rear (Fig. 4C **yellow arrows, Supplementary video 1**). This was also quantified by immunofluorescence of *CYRI-B* knockout cells (Fig. 4D-E). These C-shaped cells resembled the fast-moving goldfish keratocyte ^31^ and moved faster than cells displaying other shapes in videos. Migration speed correlated positively with the time spent as a C-shape (Fig. 4F). *CYRI-B* knockout cells showed a higher percentage of C-shaped cells when migrating, and spent a larger proportion of their time as a C-shape than wild type cells (Fig. 4G,H). Finally, the C-shaped cells migrated faster than randomly shaped cells in both wild type and *CYRI-B* knockouts, and *CYRI-B* knockouts of both C- and random shapes travelled faster than wild type counterparts (Fig. 4I). Our results suggest that *CYRI-B* knockout cells have a uniformly high state of active Rac1 and Scar/WAVE complex at their periphery, and consequently less ability to form complex multi-protrusion shapes. Cells with a uniform lamellipod all around the perimeter do not move efficiently, but if they break symmetry and assume a C-shape, the lamellipod in front is stable and cells migrate rapidly. The importance of symmetry breaking for migration has previously been described for cell fragments ^32^ and in experiments where symmetry was broken by adding inhibitors locally with a micropipette ^33^. There is an optimal level of Rac1 activity for efficient migration ^29, 34^ and in adherent cells, we propose that loss of CYRI enhances Rac1 activity and thus migration speed at the expense of complexity and plasticity of pseudopod extensions.

### CYRI promotes pseudopod splitting in *Dictyostelium*

To test conservation of CYRI function, *Dictyostelium* cells (AX3, CYRI knockout and rescue) were migrated under agarose towards folate (Fig. 5A, **Supplementary Movie 3**). *CYRI* knockout cells were rounder and displayed blunted protrusions (Fig. 5A-B, **Supplementary Movie 3)** and a decreased protrusion frequency (from ~5/min to ~2/min, Fig. 5C) and a decreased rate of pseudopod splitting (Fig. 5D). We also rescued *CYRI* knockouts with CYRI or CYRI^R155/156D^ (mutated analogously to mammalian CYRI-B^R160/161^) as stable, single-copy REMI transfectants^35^ under an actin15 promoter (Fig. 5 **A-D, Supplementary Movie 3**). CYRI^WT^ expressing cells showed increased complexity at their leading edges over wild-type cells, exhibiting more numerous fractal and branched pseudopods (Fig. 5A). As expected, CYRI^WT^ restored the circularity and enhanced the frequency of protrusion generation (around 7/min, Fig. 5C) and an enhanced rate of pseudopod splitting (around 3.5/min, Fig. 5D) even over WT cells. The enhanced rates and slightly fractal appearance of rescued cells were presumably due to mild overexpression; similar results were obtained with independent knockouts and extrachromosomal re-expression (Supplementary Fig. 5A-C). The migration speed of *CYRI* null cells was modestly reduced (from around 0.25um/sec to 0.15um/sec) and was rescued with re-expression of CYRI^WT^ but not mutant CYRI^R155/156D^ (Supplementary Fig. 5D), confirming the importance of CYRI’ interaction with Rac. The chemotactic index of cells migrating toward a folate gradient was barely affected by knockout or re-expression, remaining around 0.5-0.6 (Supplementary Fig. 5E). Pseudopod protrusion correlates with persistent migration, so we examined the directness of migration, by comparing the distance between two points and the total path length taken by cells migrating between them. Loss of CYRI caused a small increase in the directness (Fig. 5E), reflecting the increased tendency to create single broad stable pseudopods as they migrated. As expected, CYRI^WT^ rescue decreased the directness of migration in line with the fractal nature of the cells, and the mutant CYRI^R155/156D^ was not significantly different from the knockout, giving no rescue (Fig. 5E).

**Figure 5:**
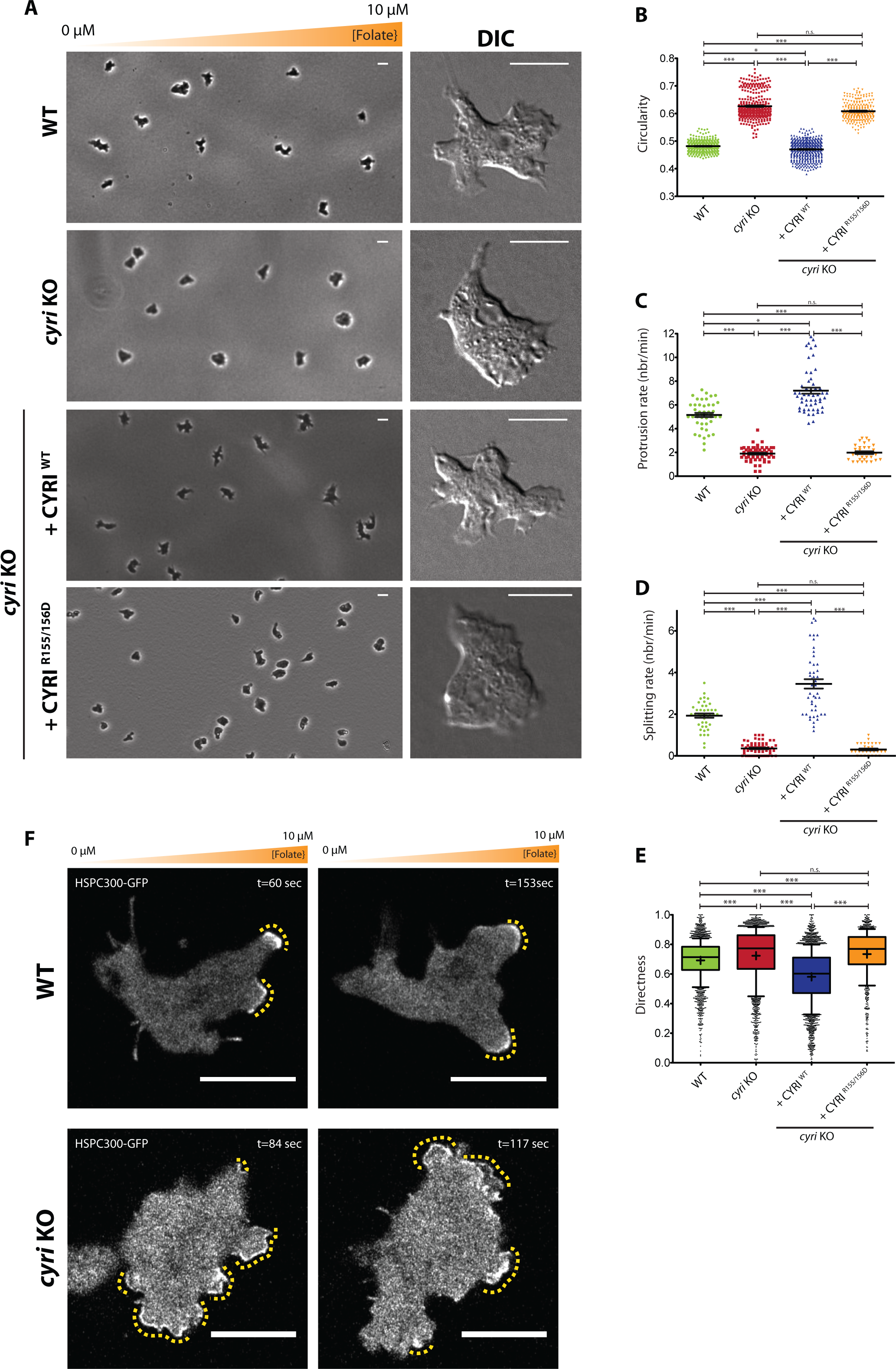
Knockout of *Dictyostelium* CYRI drives increased pseudopod splitting. A) Representative phase contrast pictures from an under agarose chemotaxis assay of Ax3 (WT) and Ax3-derived cell lines migrating towards a folate gradient, as illustrated. Representative DIC pictures from the same experiment are shown next to each panel. Cells were rescued using a restriction enzyme mediated integration (REMI) strategy (see Methods). Quantification of cell circularity (B), protrusion rate (C) splitting rate (D), directness (E) from WT, *CYRI* knockout and REMI rescued strains. See also Supplementary Movies 2 and 3. F) Still picture from a timelapse movie presenting WT and *CYRI* KO strain, transfected with HSPC300-GFP and migrating in a squashed under agarose chemotaxis assay. Yellow dotted lines highlight the length of HSCP300-GFP accumulation along the cell edge. Scale bar = 10 μm. See also Supplementary Movie 4. *B: n>240 cells per condition. One-Way ANOVA with Dunn’s test. n.s. p>0.05, *: p ≤0.05, ***: p ≤0.001* *C: n>30 cells per condition from the entire timelapse. One-Way ANOVA with Dunn’s test. n.s. p>0.05, *: p ≤0.05; ***: p ≤0.001* *D: n>30 cells per condition from the entire timelapse One-Way ANOVA with Tukey’s test. n.s. p>0.05, ***: p ≤0.001* *E: n>1200 cells per condition. One-Way ANOVA with Dunn’s test. n.s. p>0.05, ***: p ≤0.001* All data shown in this figure are representative of at least 3 independent experiments. Scale bars = 20 μm (Phase contrast images) and 10 μm (DIC images) Scatter plots and bar plot show data points with mean and SEM. Whisker plots show 10-90 percentile and mean (cross).

As in mammalian cells, altering CYRI affected recruitment of Scar/WAVE complex to the edges of migrating cells, leading to broad pseudopods and altered motility. The Scar/WAVE complex is recruited to the pseudopods of both WT and *CYRI* knockout cells with a very different profile (Fig. 5F **and Supplementary Movie 4**). Scar/WAVE complex (indicated by HSPC300-GFP ^5^) forms a thick and restricted patch at the ends of extended pseudopods whereas the signal is more noisy, diffuse and broad in the knockout cells. This phenotype is similar to mammalian *CYRI-B* knockout cells, where extensive Scar/WAVE2 recruitment was apparent around the cell periphery in broad regions. Thus, CYRI has a conserved function to restrict the accumulation of the Scar/WAVE complex at the cell edge and to sharpen and enhance individual pseudopods to provide control of cell migration.

### CYRI is a locally acting negative inhibitor of actin assembly signals

It is widely accepted that actin assembly pathways are not linear cascades, but rather feedback loops where positive stimulation is self-reinforcing and causes further activation until overcome by negative feedback^36, 37^. In models of migration based around positive feedback, for example the mathematical model of Meinhardt ^38^, inhibitors are required to limit the amount of cell edge devoted to pseudopods. A locally-acting inhibitor is also needed to destabilise existing pseudopods, so the cell can change direction. Thus the effect of the local inhibitor is to make pseudopods more dynamic, with old pseudopods being removed, and new ones being created. We used a published simulation^39^ based on the Meinhardt model^38^ to visualise the concentrations of the activator and the local inhibitor at the cell edge (Supplementary Fig. 6 **and Supplementary Movie 5**), to provide a clear illustration of the proposed role for CYRI-B in regulation of Rac1 and Scar/WAVE signaling. A peak in the activator (which represents, for example, active Rac and Scar/WAVE) results in the formation of a new pseudopod. The peak also causes an increase in the concentration of the local inhibitor, which is smaller and thus diffuses faster. This causes two changes – initially, it limits the lateral spread of the pseudopod (Supplementary Fig. 6, panel 1); later, levels of inhibitor rise in the middle of the pseudopod, destabilizing it and causing it to split (Supplementary Fig. 6, panel 2). The weaker of the pseudopods then retracts and the stronger is reinforced until the cycle of inhibition catches up with it and starts the splitting all over again (Supplementary Fig. 6, panels 3-4). The local inhibitor thus increases both the morphological complexity of the cell and the competition between pseudopods. CYRI resembles Meinhardt’s local inhibitor in several ways. Its loss causes pseudopods to spread; overexpression makes pseudopods in both mammalian and *Dictyostelium* cells narrower. Similarly, pseudopods in the cells lacking CYRI are far less dynamic, while they change rapidly in overexpressers. Thus, Meinhardt’s model offers insight into the role of CYRI proteins as local inhibitors which perhaps counterintuitively enhance leading edge dynamics and add plasticity to the positive feedback loops driving migration.

## Discussion

The broader implications of CYRI modulation of actin cytoskeletal control remain to be determined for normal and tumour cells. Modulating CYRI expression levels can either speed up or slow down cell migration depending on the circumstances. For example, we saw faster migration in 2D assays (Fig. 4) and increases in 3D invasion in CHL1 cells (Supplementary Fig. 4) but *Dictyostelium* cells lacking CYRI migrated modestly more slowly (Supplementary Fig. 5D). Migration speed is a complex parameter and depends on a balance between many factors including adhesion, protrusion and contractility. *Dictyostelium* cells already show optimal migration due to constitutive front-rear polarization and relatively low adhesion to rigid surfaces, so a change in the balance of Rac1 and Scar/WAVE activation may slow them down. Mammalian cancer cells or fibroblasts migrate orders of magnitude more slowly on similar surfaces, so increases in Rac1 signaling upon loss of CYRI-B may thus increase their speed. The gene encoding CYRI-B is located on chromosome 8q24, near Myc and is thus genetically co-amplified with the Myc oncogene in many cancers (C Bioportal database). Coupled with a potential role in invasion, which we highlight here (Supplementary Fig. 4), we predict an interesting role for CYRI in cancer.

CYRI proteins represent new fundamental direct negative regulators of the Rac1 – Scar/WAVE pathway and the DUF1394 is here identified as a new module specifically binding to active Rac. Previous studies have identified negative regulation at the level of Arp2/3 complex and positive regulators of Scar/WAVE complex such as lamellipodin ^40, 41^, but CYRI is the only locally acting negative regulator of Rac1-Scar/WAVE activation. CYRI competes with CYFIP to interact with active Rac1 and thus dampens the activation of Scar/WAVE complex by active Rac1. This local dampening of the signal focuses activation to the central regions of the highest active zones and thus sharpens pseudopod protrusions. CYRI thus allows the cell to generate highly flexible and dynamic protrusions that can split and facilitate changes in direction. How CYRI is regulated and the structural nature of its interaction with Rac1 are among the many interesting questions for the future.

## Acknowledgements

We thank Margaret O’Prey and the BAIR imaging facility for help with microscopy, Chloe Tesniere for careful work with CYRI overexpression plasmids, Michael Mcilwraith for help with protein purification, Luke Tweedy for helpful advice on quantification and image analysis, Benjamin Tyrell for isolation of the inducible Rac ^fl/fl^ fibroblasts and David Bryant for advice and constructs. We thank CRUK for core funding to L.M.M. (grant A15673), R.H.I. (grant A19257) and S.Z. (C596/A12935), and BBSRC for funding to L.H.C and N.C.O.T (BB/L022087/1).

## Author contributions

R.H.I. and J.B. conceived and carried out the initial screen, the *Dictyostelium* experiments and recognized the similarity of CYRI to CYFIP. L.F. designed and carried out the majority of the experiments on mammalian CYRI-B. L.M.M., R.H.I. and L.F. conceived the study and wrote the paper. K.M. and K.I.A. designed the Raichu FRET probe and with L.F. carried out the FRET experiments. P.B. and L.F. carried out the surface plasmon resonance experiments. J.G., N.C.O.T., and L.C. synthesized probes for, advised on and carried out the myristoylation experiments with L.F. H.J.S. provided technical help and carried out experiments. S.L. and S.Z. carried out and analysed the mass spectrometry with L.F. and J.B. P.T. and S.I. provided essential advice and carried out experiments and analysis of data. M.N. and R.H.I. constructed the model and advised on its use.

## Supplementary Movie 1

Random migration assay of control or CYRI-B CrispR knockout CHL1 cells, plated on fibronectin and imaged every 10 minutes. Movies are representative of a 8h timelapse experiment.

Scale bar = 100 μm

## Supplementary Movie 2

Tiles phase contrast movies from a under agarose chemotaxis assays of cells chemotaxing toward folate, with each strain as indicated.

Scale bar = 50 μm 1frame/min

## Supplementary Movie 3

Tiled DIC movies from under agarose chemotaxis assays of WT, *CYRI* KO, or the REMI rescue strains (*CYRI* KO + WT CYRI and *CYRI* KO + R155/156D CYRI, as indicated) migrating toward folate.

1 frame/2sec Scale bar = 5 μm

## Supplementary Movie 4

Under agarose chemotaxic assay toward folate of wild type or *CYRI* KO Ax3 cells transfected with HSPC300-GFP.

Pictures were taken every 3 seconds. Scale bar = 10 μm.

## Supplementary Movie 5

Simulation of a cell protrusion and motility, based on a mathematical model^5, 6^. Cell protrusions are generated following an increase of an activator (Red: Rac1 or Scar/WAVE complex), cross-talking to a local inhibitor (Blue: CYRI) that restricts pseudopod width. Accumulation of the local inhibitor causes pseudopod spitting. Time (t) is in arbitrary units.

## Supplementary Materials and Methods

**Supplementary Figure 1:**
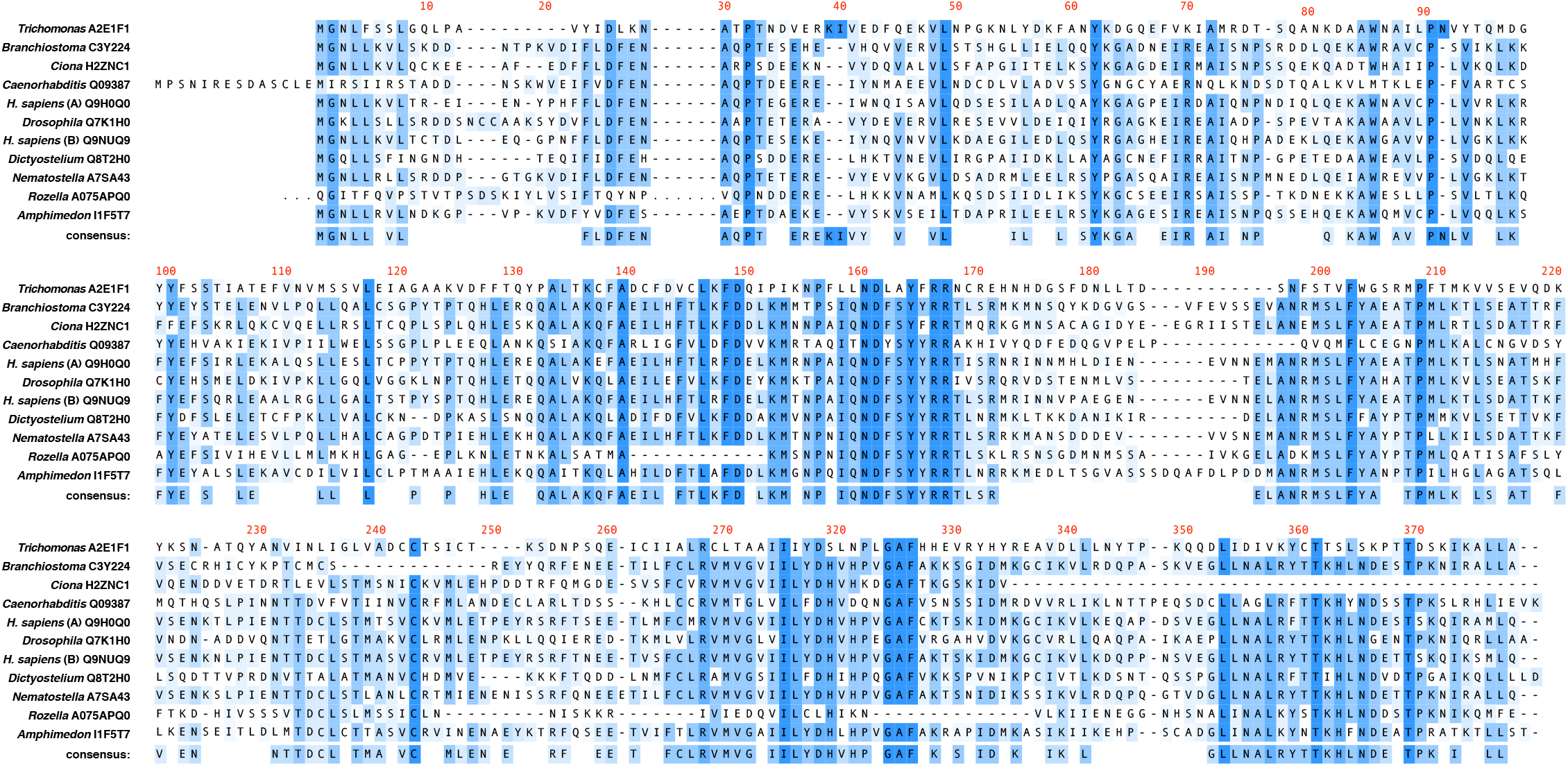
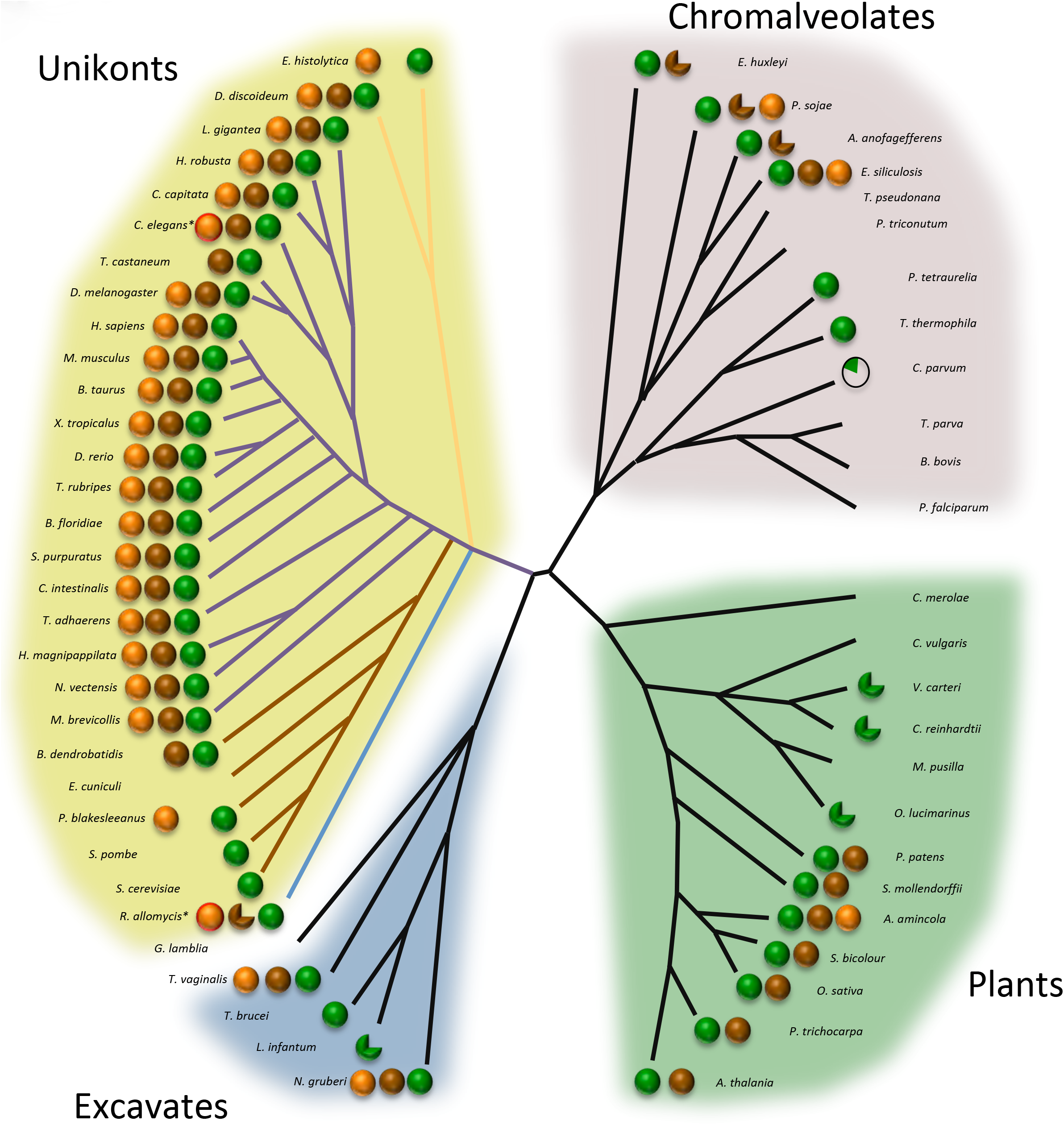
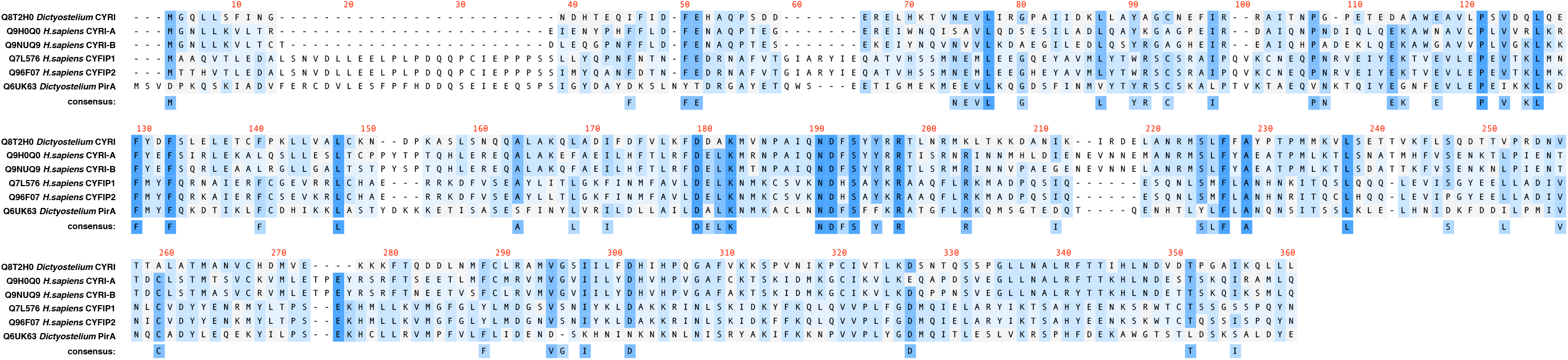
Sequence conservation of CYFIP and CYRI across the eukaryotic evolutionary tree. A)Alignment of CYRI (FAM49) from diverse representative organisms of the eukaryotic evolutionary tree. UniProt accession numbers are reported. Color code represents the number of entries with identical amino acid at each position. B)Co-evolution of Arp2/3 (green), Scar/WAVE (brown) and CYRI (orange) across the 4 main superfamilies of the eukaryotic tree ^1^. An incomplete circle corresponds to the absence of some subunits from the complex as in ^2^. Each supergroup was assigned to a specific background color: Unikonts (yellow), Chromalveolates (pink), Excavates (blue) and Plants (green). Branch length does not reflect evolutionary distances. For *C. elegans* and *R. allomycis*, * and red highlight around circle denotes lack of putative N-terminal myristoylation site on CYRI. Different coloured branches in Unikonts represent amoebazoa (yellow branches), metazoan (purple branches), fungi (brown branches) and Rozella allomycis, the earliest diverged of the fungi (blue branch). C)Alignment of *Dictyostelium discoideum* and *Homo sapiens* CYFIP and CYRI (FAM49) sequences. UniProt accession numbers are reported. Color code represents the number of entries with identical amino acid at this position.

**Supplementary Figure 2:**
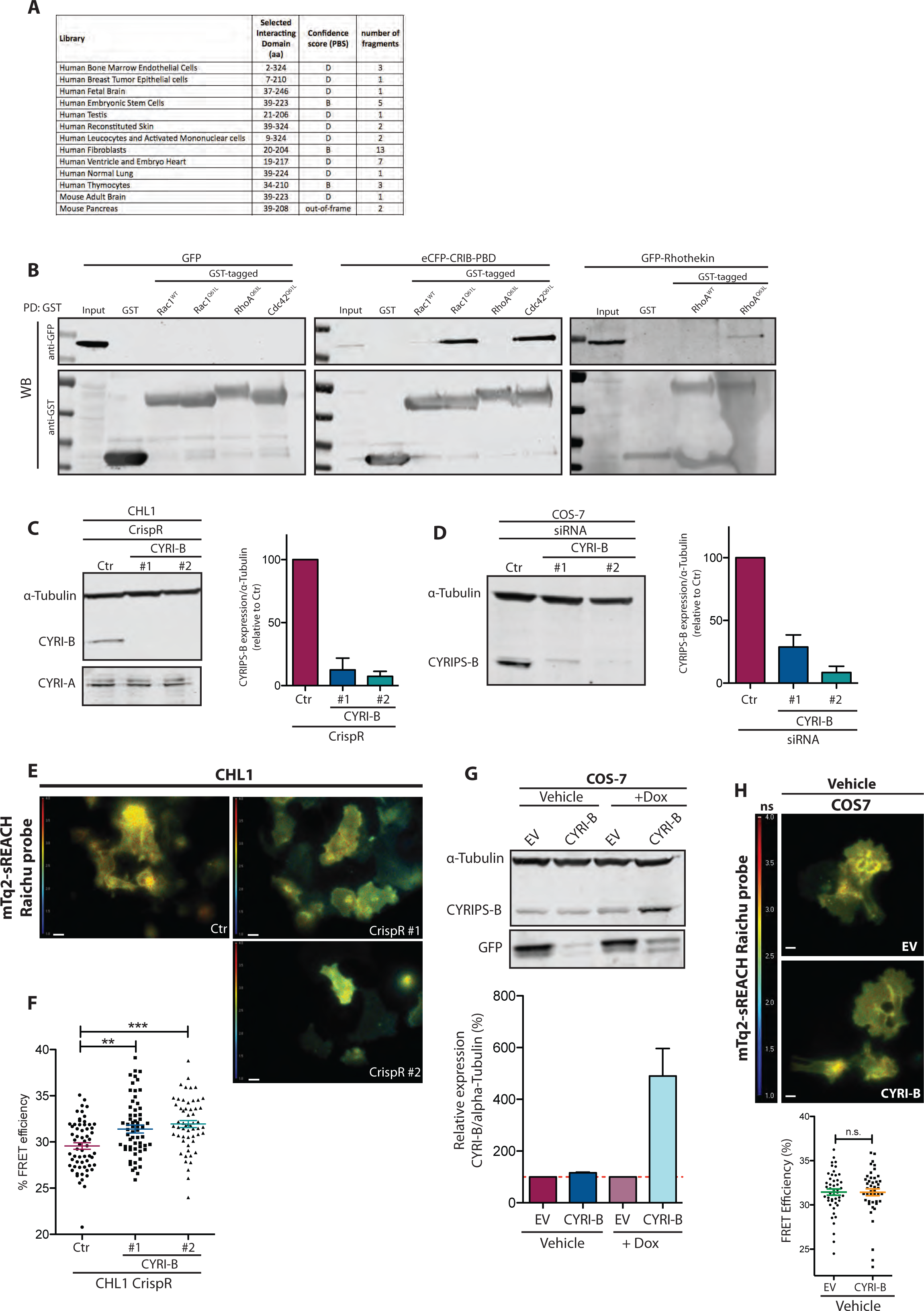
CYRI-B interacts with active Rac1 and buffers Rac1 activity at the membrane. A)Table presenting the interacting regions between CYRI-B and constitutively active Rac1^G12V^. Number of interacting fragments obtained by yeast-two hybrid is shown as well as the mapping of this region in the designated library. Confidence score explained in methods (A=best to D=low). B)Control western blots from GST-pulldowns showing specific binding to Rac1 over the other Rho GTPases. Lysate from CHL1 cells expressing GFP (negative control) or either eCFP-Crib-PBD or GFP-Rhotekin (reporter for Rac1 or RhoA activity respectively) were mixed with bacterially expressed GST, GST-Rac1^WT^, GST-Rac1^Q61L^, GST-RhoA^Q63L^, GST-Cdc42^Q61L^ and immobilized on beads. C)Establishment of 2 independent *CYRI-B* knockout CHL1 cells using CrispR-Cas9 was confirmed by western blot using 2 different guide RNA sequences #1 and #2. CYRI-B expression was quantified after puromycin selection and normalized to the control CrispR cell line. CYRI-A expression was also checked to confirm the specificity of the guide RNA. D Western blot analysis of CYRI-B expression after knockdown by siRNA in COS-7 cells. Quantification was normalized to the control scramble siRNA. E-F) FLIM imaging and quantification of control or *CYRI-b* knockout CHL1 cells The jet2 color code shows the average lifetime of the mTq2-sREACH Raichu probe, oscillating between 1ns-4ns (blue-red). *(Ctr n=61, CrispR#1 n=63, CrispR #2 n=56). Dunn’s test was performed. ** p≤0.01, *** p≤0.001.* G) Western blot analysis of COS7 cells transiently transfected with a mVenus-TetON inducible CYRI-B construct. Anti-GFP was used to probe for mVenus co-expression. Cells were collected and analysed 48h after vehicle or doxycycline treatment. Relative CYRI-B expression was normalized to the control condition (EV construct level highlighted with the red dotted line). H)FLIM imaging and quantification of COS-7 cells transfected with the empty inducible vector backbone (EV) or containing a CYRI-B cDNA (CYRI-B) and treated with vehicle solution for 48h. The jet2 colour code shows the average lifetime of the mTq2-sREACH Raichu probe, spanning 1-4ns (blue to red). *(Vehicle_EV = 47 cells, Vehicle_CYRI-B = 46 cells). Mann Whitney test was performed. n.s. >0.05.* All data shown in this figure are representative of at least 3 independent experiments. Scatter and bar plots show data points with mean and SEM. Scale bar = 10μm

**Supplementary Figure 3:**
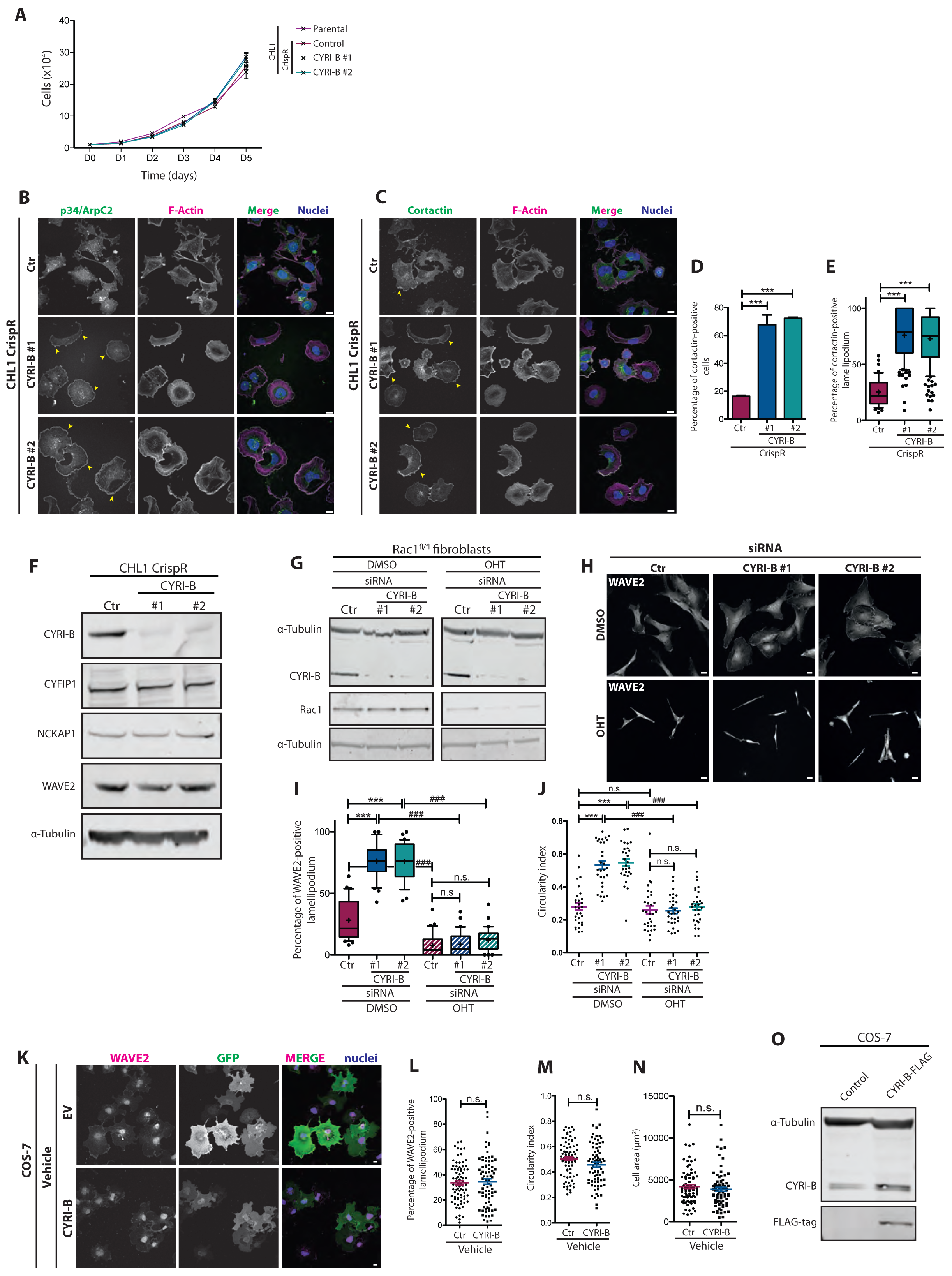
CYRI-B restrains cell protrusion formation. A) Growth curves of parental, control or *CYRI-B* CrispR knockout CHL1 cells. Cells were plated in triplicate and cell number was averaged for each repeat. B-E) Localisation of endogenous p34/ArpC2 (green) (B), or Cortactin (green) (C) and F-actin (magenta) in control (Ctr) or *CYRI-B* CrispR knockout cells, plated 4h on collagen. Arrowheads show the accumulation of p34/ARPC2 and Cortactin at the lamellipodium. Quantification of percentage of cells showing a cortactin staining at the lamellipodium (D) and extent of the cortactin staining over the total cell perimeter (E). *Dunn’s test *** p≤0.001* *(Ctr n=44, CrispR#1 n=145, CrispR #2 n=152)* F)Western blot analysis of control or *CYRI-b* CrispR knockout CHL1 cells, probe for Scar/WAVE complex members and alpha-Tubulin. G)Western blot analysis of ROSA26::Cre-ER^T2+^;p16Ink4a^−/−^; Rac1^fl/fl^ mouse tail fibroblasts treated 7 days with DMSO or OHT to induce Rac1 recombination and knockdown for CYRI-B by siRNA. H)Immunfluorescence pictures of control or Rac1 knockout rat tail fibroblasts treated with control or siCYRI-B, plated on fibronectin and stained for WAVE2. Quantification of the WAVE2 staining at the lamellipodium and circularity are displayed in I and J respectively. One-Way ANOVA with Dunn’s test *(n.s p>0.05,* *\*\*\* p≤0.001*) or Mann Whitney t-test (*n.s p>0.05, ### p≤0.001*). *(30 cells/condition)* K) Immunofluorescence of COS-7 cells transiently transfected with the inducible mVenus-TetON construct and treated with the vehicle solution. Cells were allowed to spread on fibronectin for 4h and stained for endogenous WAVE2 and GFP. Only GFP expressing cells were used for further quantification. Percentage of lamellipodium covered by a WAVE2 staining (L), cell circularity (M) and cell area (N) were quantified. *Mann Whitney t-test was performed. n.s p>0.05. (Ctr=81 cells, CYRI-B=78cells)* O) Western blot from wt or CYRI-B-FLAG transfected COS-7 cells. All data shown in this figure are representative of at least 3 independent experiments. Scatter plots and bar plots show data points with mean and SEM. Whisker plots show 10-90 percentile and mean (cross). Scale bar = 10μm.

**Supplementary Figure 4:**
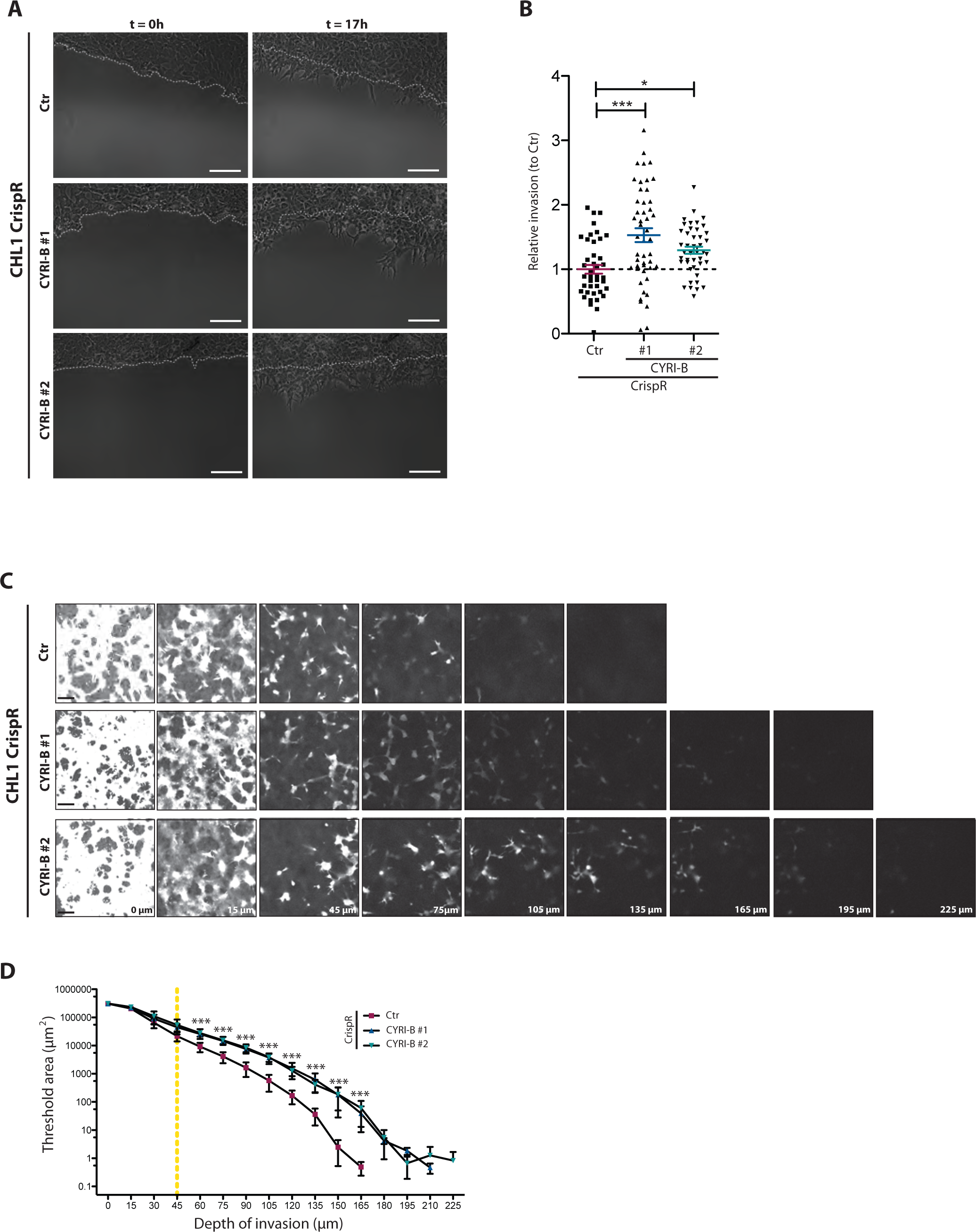
CYRI-B negatively regulates 2D and 3D cell invasion. A) Circular invasion assay^3^ of control or *CYRI-B* CrispR knockout CHL1 cells invading through Matrigel. White dotted lines indicate the start of the invasive front at t=0h. Invasion index after 17h is plotted in B for each movie. *(Ctr n=43 movies, CrispR#1 n=48 movies, CrispR#2 n=45 movies).* One-Way ANOVA with Dunn’s test *\*: p ≤0.05; ***: p ≤0.001* C) Inverted Invasion Assay^4^ of Control (Ctr) or *CYRI-B* CrispR knockout CHL1 cells. Cells were allowed to migrate through a 8 μm pore filter and invade through a Matrigel plug above the filter. Serial optical sections were captured at 15 μm intervals and presented as a sequence in which the individual optical sections are placed alongside one another with increasing depth from left to right as indicated. Images at 0 μm indicate cells that came though the filter but did not enter the gel. Invasion was plotted in D for each optical section throughout the whole matrigel plug using a calcein staining as a cell marker. Cells migrating above 45 μm of depth (dotted line) are considered as invasive. 5 fields across the filter of each technical replicate were taken for each experiment. Likelihood ratio test with Bonferroni correction test. *** p<0.001 All data shown in this figure are representative of at least 3 independent experiments. Error bars represent S.E.M Scale bar = 100 μm.

**Supplementary Figure 5:**
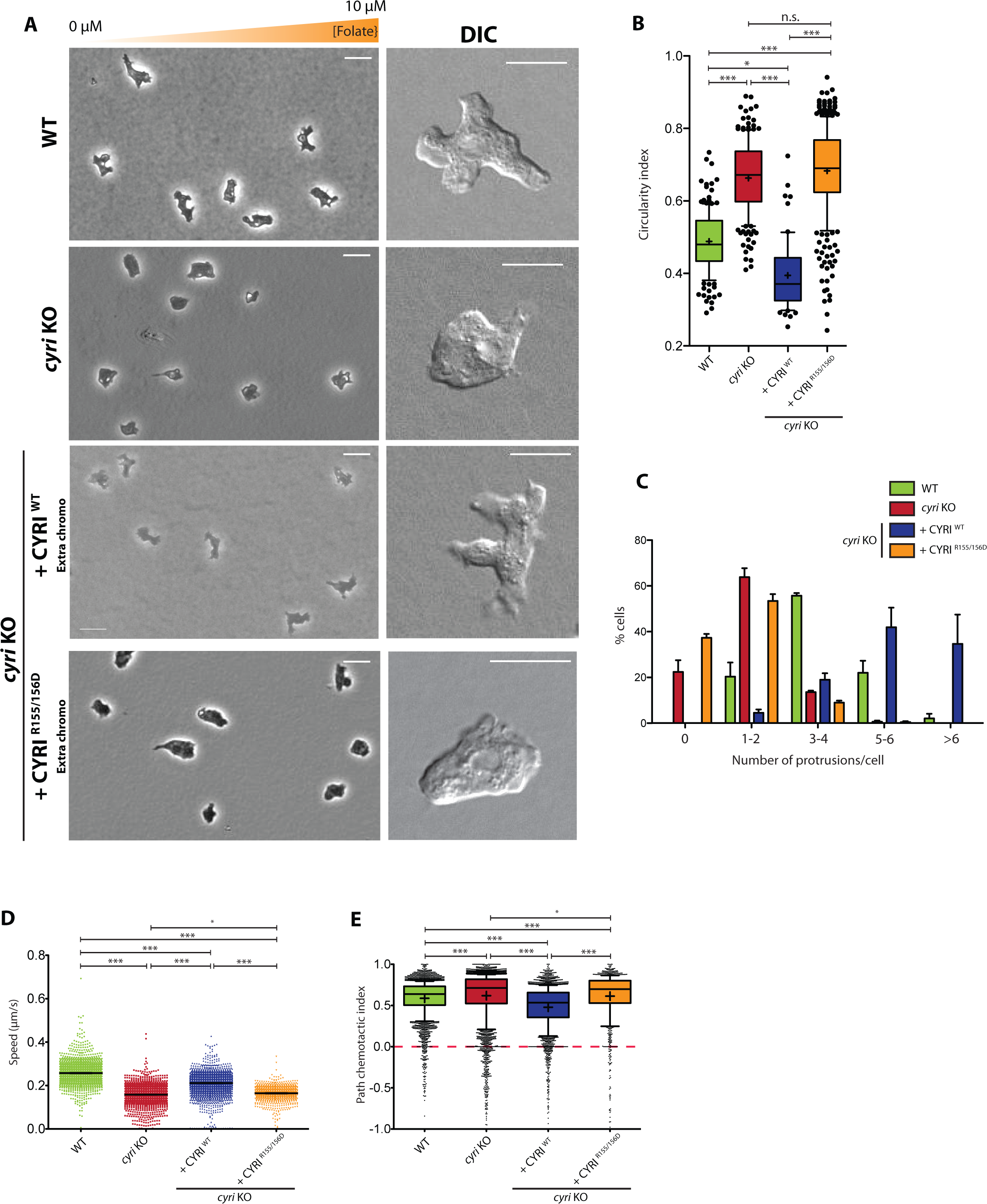
Effect of CYRI in *Dictyostelium* morphology. A) Similar to Figure 5, this figure shows the effect of CYRI KO rescue with CYRI cloned into a multi-copy extrachromosomal plasmid. Representative phase contrast pictures from an under agarose chemotaxis assay of Ax3 (WT) and Ax3-derived cell lines migrating towards a folate gradient. DIC images from the same experiment are shown next to each panel. B) Cell circularity values were calculated from cell outlines drawn manually on stills. Number of protrusions per cell was manually quantified (C) using random fields from movies of the under agarose chemotaxis assay. Automatic quantification of the cell speed (D) and path chemotactic index (E) from WT, *CYRI* knockout and REMI rescue strains. Shown in Figure 5. D-E: n>1200 cells per condition. One-Way ANOVA with Dunn’s test. n.s. *p*>0.05, *\*: p ≤0.05; ***: p ≤0.001* All data shown in this figure are representative of at least 3 independent experiments. Scale bars = 20 μm (Phase contrast images) and 10 μm (DIC images). Scatter plots and bar plot show data points with mean and SEM. Whisker plots show 10-90 percentile and mean (cross).

**Supplementary Figure 6:**
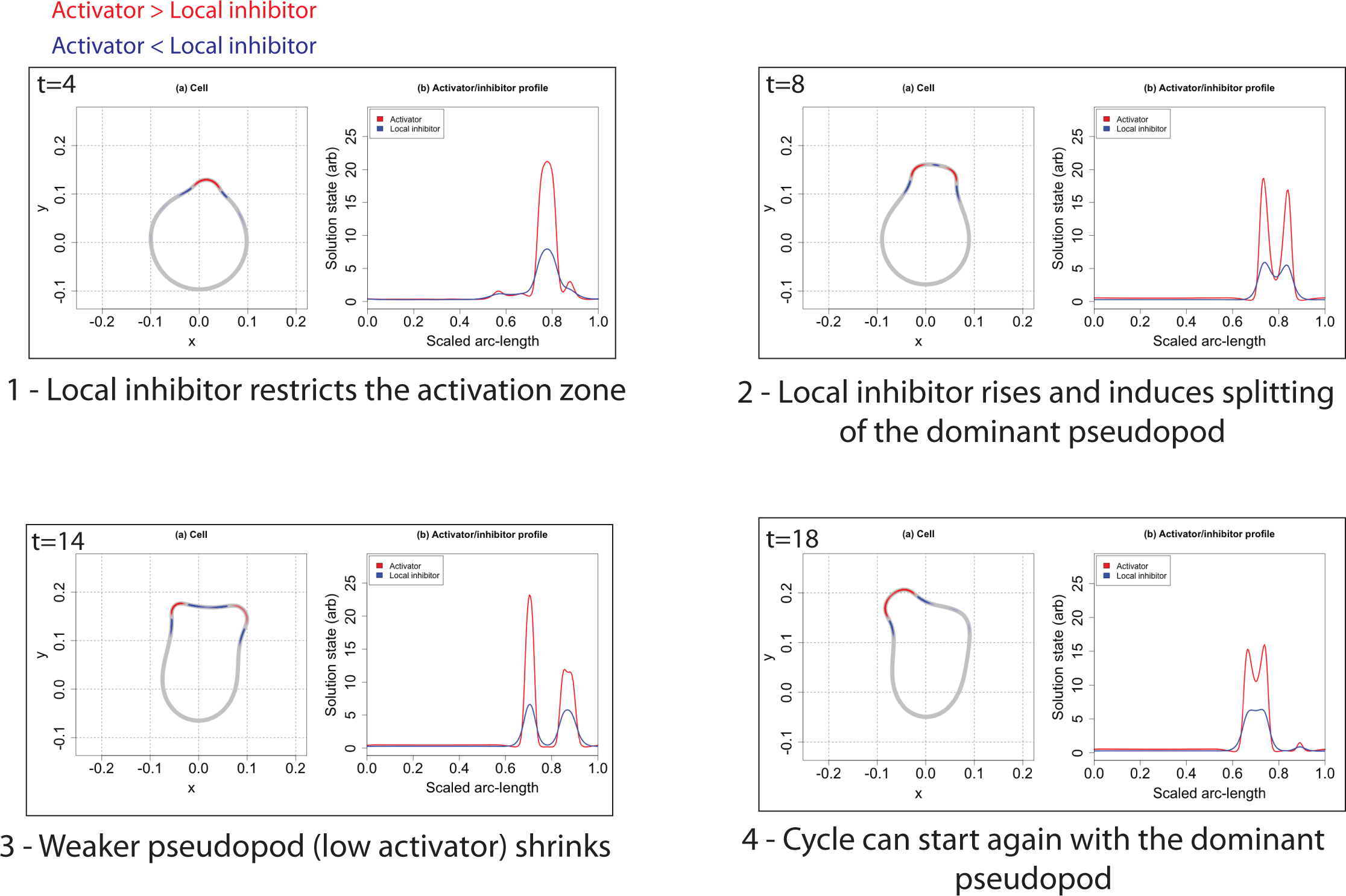
Model depicting how a local inhibitor affects pseudopod formation and complexity. In each panel, graph (a) shows the outline of a simulated cell, with red areas of the periphery having more activator than local inhibitor and blue having more local inhibitor than activator. The x- and y-axes correspond to arbitrary units representing distance. The simulation was run from time t=0 and panels are taken from times shown (arbitrary units of time). Graph (b) shows the concentrations of the activator (red) and the local inhibitor (blue), where the x-axis represents the scaled arc-length around the perimeter of the cell and the y-axis represents the concentration in arbitrary units. Panel 1 (t=4) shows the generation of a pseudopod, where the local inhibitor causes the edges of the activated region to sharpen. Panel 2 (t=8) shows a split in the activator profile (and, consequently, a split in the associated pseudopod), which results from the higher relative concentration of the local inhibitor near the centre of the activated region. Panel 3 (t=14) shows the cell with two smaller pseudopods, which compete for dominance and determine the direction of migration. Panel 4 (t=18) shows the winning pseudopod starting the cycle over again and the local inhibitor rising in response to the rise in activator.

**Table S1:**
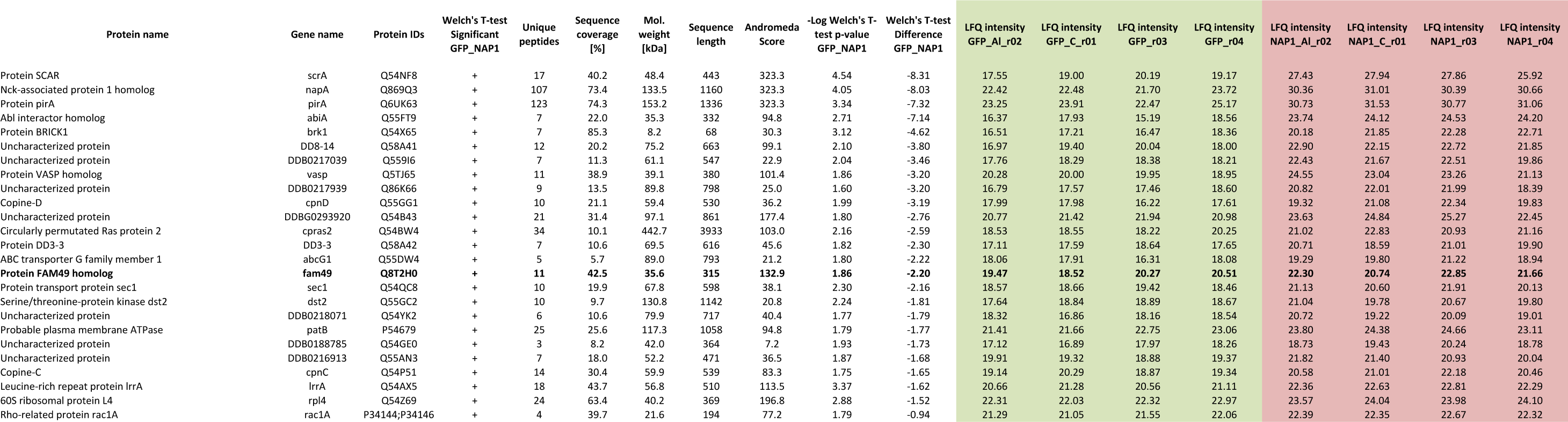
Significant candidates obtained from Nap1 pull down after reversible crosslink treatment. Uniprot accession numbers are mentionned for each putative interactor. Welch’s t test and further analysis were performed using the MaxQuant software.

**Table S2:** Antibody List

| **Antibody** | **Source** | **Dilution** |
| --- | --- | --- |
| Rabbit anti Fam49B | Sigma #HPA009076 | WB 1:500 |
| Rabbit anti Fam49B | ProteinTech #20127-1-AP | WB 1:1000 |
| Mouse anti GFP (4B10) | Cell Signaling #2955 | WB 1:1000 |
| Mouse anti GFP (JL-8) | ClonTech #632380 | WB 1:1000 |
| Rabbit anti GFP | Abcam #ab290 | WB 1:2000 |
| Chicken anti GFP | Abcam #ab13970 | IF 1:500 |
| Mouse anti GST | Abcam #ab18183 | WB 1:1000 |
| Rabbit anti GST | Cell Signaling #2622 | WB 1:1000 |
| Rabbit anti WAVE2 (H-110) | SantaCruz #sc-33548 | IF 1:50<br>WB 1:200 |
| Mouse anti FLAG-tag (M2) | Sigma #1804 | IF 1:400 |
| Mouse anti FLAG-tag (9A3) | Cell Signaling #8146 | WB 1:1000 |
| Mouse anti Cortactin (4F11) | Millipore #05-180 | IF 1:200 |
| Rabbit anti p34-Arc/ARPC2 | Millipore #07-227 | IF 1:200 |
| Mouse anti Tubulin (DM1A) | Sigma #T9026 | WB 1:3000 |
| Rabbit anti Fam49A | Sigma #SAB1103179 | WB 1:1000 |
| Rabbit anti Cyfip1 | Abcam #156016 | WB 1:1000 |
| Rabbit anti p125Nap1 | MilliPore #07-515 | WB 1:1000 |
| Rabbit Rac1/2/3 | Cell Signaling #2465 | WB 1:1000 |
| Goat anti Rabbit 800nm | Thermo Scientific #SA5-35571 | WB 1:10000 |
| Goat anti Mouse 800nm | Thermo Scientific #SA5-35521 | WB 1:10000 |
| Donkey anti Rabbit 680nm | Invitrogen #A21206 | WB 1:10000 |
| Donkey anti Mouse 680nm | Invitrogen #A10038 | WB 1:10000 |
| Donkey anti Mouse 594nm | Invitrogen #A31203 | IF 1:200 |
| Goat anti Rabbit 488nm | Invitrogen #A11008 | IF 1:200 |
| Goat anti Mouse 488nm | Invitrogen #A11001 | IF 1:200 |
| Donkey anti Chicken 488nm | Invitrogen #A11039 | IF 1:200 |
| Phalloidin 488nm | Molecular Probe #A12379 | IF 1:200 |
| Phalloidin 594nm | Molecular Probe #A12382 | IF 1:200 |
| Hoechst 33342 | Thermo Scientific #62249 | IF 1:50000 |

### Reagents

All chemicals were purchased from Sigma-Aldrich unless specified otherwise. Antibodies are listed in Table S2 below.

### Alignment and phylogenetic tree

Protein sequences were obtained from Uniprot http://www.uniprot.org/ and aligned using MacVector software. The phylogenetic tree was constructed based on the major eukaryotic superclasses as previously defined ^1^ and based on previous identification of Arp2/3 complex and Scar/WAVE complex sequences in selected organisms representing branches of this tree ^2, 7^. Searches were performed for the presence of CYRI using a BLAST homology search on the NCBI website <u>https://blast.ncbi.nlm.nih.gov/Blast.cgi</u>. *Dictyostelium* or human or a closer relative from the tree were searched against the complete translated genome of open reading frames from these organisms.

HMM logo was generated by feeding the Pfam seed database of the DUF1394 domain into Skylign^8^.

### Mammalian cell lines and growth conditions

CHL1, B16F1, HEK293T, COS-7 cells were maintained in 10% FBS and 2mM L-Glutamine supplemented Dulbecco’s Modified Eagle’s Medium. ROSA26:CreER^t2+;^ Ink4^−/−^; Rac^fl/fl^ mouse tail fibroblasts were maintained in the aforementioned medium complemented with 1mg/mL of primocyn. Recombination was obtained by adding 1μM of hydroxytamoxifen during 7 days.

COS-7 cells transfected with the Doxycycline inducible system were grown in 10% of tetracyclin-free FBS (ClonTech) and treated with 5μg/mL of doxycycline for 48h. All mammalian cell lines used in this study were maintained in ø10cm plastic dishes at 37 °C and perfused with 5% CO_2_.

### CLICK Chemistry of Mammalian CYRI-B

HEK293T cells plated on 24-well plate were transfected with 1 μg of pEGFPN1 or CYRI-B-EGFP (wild-type or G2A mutant) using Lipofectamine 2000 reagent and were processed the next day. C14:0-azide was synthesised as previously described ^9^. Transfected HEK293T cells were incubated with 100 μM of C14:0-azide (in DMEM with 1 mg/mL defatted BSA) for 4 h at 37 °C. Cells were washed twice in PBS and lysed on ice for 10 min in 100μL lysis buffer (150mM NaCl, 1 % Triton X-100, 50mM Tris-HCl, pH8.0, containing protease inhibitors). Cell lysates were centrifuged at 10, 000 x g for 10 minutes at 4 °C to remove cell debris.

Conjugation of alkyne IR-800 Dye to C14:0 azide was carried out for 1 h at room temperature with end-over-end rotation by adding an equal volume of freshly mixed click chemistry reaction mixture containing 10 μM 800CW alkyne infrared dye, 4 mM CuSO4, 400 μM Tris[(1-benzyl-1H-1,2,3-triazol-4-yl)methyl]amine, and 8 mM ascorbic acid in dH_2_0 to the supernatants. GFP-tagged proteins were isolated using the μMACS GFP isolation kit following manufacturer’s protocol and resolved by SDS-PAGE as previously described. Protein acylation was quantified by expressing the intensity of the CLICK signal relative to the protein signal.

### Yeast Two-Hybrid screen

Screening was performed at Hybrigenics services *as per* their standard protocols. Briefly, the coding sequence for the constitutively active full-length Rac1 (NM_006908.4; mutations G12V, C189S) was PCR-amplified and cloned into pB27 as a C-terminal fusion to LexA (LexA-Rac1). All libraries that were used to screen are domain libraries cloned into the prey vector pP6. pB27 and pP6 are derived from the original pBTM116 ^10^ and pGADGH ^11^ plasmids, respectively.

The bait was screened against the different libraries using a mating approach with YHGX13 (Y187 ade2-101::loxP-kanMX-loxP, mat alpha) and L40deltaGal4 (mat-a) yeast strains as previously described ^12^. Positive colonies were selected on a medium lacking tryptophan, leucine and histidine, and supplemented with 3-aminotriazole whenever necessary to reduce the total number of clones. The prey fragments of the positive clones were amplified by PCR and sequenced at their 5’ and 3’ junctions. The resulting sequences were used to identify the corresponding interacting proteins in the GenBank database (NCBI) using a fully automated procedure. A confidence score (PBS, for Predicted Biological Score) was attributed to each interaction as previously described ^13^.

### GST Pull-down of Mammalian CYRI-B and GTPases

DH5alpha *E. coli* cells were grown at OD_600nm_ 0.4 and induced for 4h with 0.2mM IPTG. Pellet was resuspended in ice-cold buffer A (50mM NaCl, 50mM Tris-HCl pH7.5, 5mM MgCl_2_, 3mM DTT) and sonicated, followed by a 30min spin at 20000 rpm to yield lysate. GST tagged proteins were immobilized 30min at 4°C with gentle agitation on pre-washed glutathione-sepharose beads and unbound proteins were washed out 3 times in buffer A.

CHL1 cells transfected with GFP constructs were collected in ice-cold lysis buffer (100mM NaCl, 25mM Tris-HCl pH7.5, 5mM MgCl_2_, 1x protease and phosphatase inhibitors, 0.5% NP-40). 2mg of proteins were mixed with pre-equilibrated beads and agitated 2h at 4°C. Beads were then washed 3 times in washing buffer (100mM NaCl, 25mM Tris-HCl pH7.5, 5mM MgCl_2_), resuspended in sample buffer containing DTT and resolved by SDS-PAGE and blotted as described below.

### Mutagenesis of Mammalian CYRI-B

Point mutation (lower case) was inserted using the Q5-site directed kit (New England Biolabs) and following the manufacturer’s instructions. Primers were designed using NEBaseChanger.

R160D (Fwd: 5’ – CAGCTATTATgacAGAACATTGAGTCGTATG-3’ Rev: 5’ – AAATCATTCTGTATGGCAGG-3’)

R161D (Fwd: 5’ – CTATTATAGAgacACATTGAGTCGTATGAGG-3’ Rev: 5’ – CTGAAATCATTCTGTATGGC-3’)

R160/161D (Fwd: 5’ - CAGCTATTATgacgacACATTGAGTCGTATGAGGATTAAC-3’ Rev: 5’ – AAATCATTCTGTATGGCAGG-3’)

### Protein purification for SPR analysis

*E. Coli* BL21 CodonPlus (DE3)-RIL (Agilent Tech.) and *E. Coli* BL21 (DE3) pLysS (Promega) were used for GST-tagged and His-Tagged proteins respectively. Pre-culture was grown overnight in L-Broth (LB) containing appropriate antibiotics. Once reaching OD_600nm_ 0.4, protein expression was induced using

0.2mM IPTG and culture was kept overnight at 20°C, 200 rpm. Cells were lysed in Buffer 1 (200mM NaCl, 30mM Tris pH7.5, 5mM MgCl_2_, 3mM β-mercaptoethanol) containing protease inhibitors (Roche) and passed through a 20000psi-pressurised microfluidizer. The soluble fraction was collected by centrifugation and loaded onto an equilibrated GSTrap HP or HisTrap HP column using an AKTA machine. Proteins were either directly eluted using Buffer 1 containing 20mM GSH (GST protein)/300mM Imidazol pH7.5 (His protein) or cleaved overnight on the column.

Proteins were gel purified (HiLoad 16/600 Superdrex 75pg or HiLoad 16/600 Superdrex 200pg) in Buffer 2 (150mM NaCl, 25nM Tris-HCl pH7.5, 5mM MgCl_2_, 2mM β-ME), snap-frozen and stored at -80°C.

### Surface Plasmon Resonance (SPR) protein binding assay

SPR analysis was performed using Biacore T200 (GE Heathlcare). Using GF buffer (200mM NaCl, 30mM Tris-HCl pH7.5, 3mM beta-mercaptoethanol) complemented with 5mM MgCl_2_ and 0.5% of surfactant P20 as running buffer, GST-tagged proteins were immobilised at 22°C onto CM5 sensor chip functionalized with anti-GSTand reached ~320 RU. Same procedure was used for His-tagged protein onto NTA sensor chip and reached 650 RU. All immobilisation steps were done at a flow rate of 10μL/min. Serial dilution of each analyte was injected across a reference flow cell and the flow cell containing the ligand at a flow rate of 30μL/min. Data were solvent corrected, reference subtracted, quality controlled and evaluated using the Biacore T200 evaluation software. Affinity was determined by curve fitting a 1:1 binding model.

### Transfection, siRNA Treatment and Knockout Mammalian Cells

Cells were plated a day before transfection at 70% of confluence and later transfected using Lipofectamine 2000 according to the manufacturer’s instructions. siRNA oligonucleotides targeting CYRI-B were purchased from Qiagen: 75 nM of Mouse CYRI-B siRNA – siRNA #1: 5’-CAGGCTCTCGCTAAACAGTTT-3’, siRNA #2: 5’-TCAGGTGAATGTAGTGTTAAA-3’ and 25nM of Monkey CYRI-B siRNA #1: 5’ – AGGGTAATGGTGGGTGTCATA-3’, siRNA #2: 5’-ATAGAAGAACATTGAGTCGTA-3’or matched concentration of control siRNA (AllStars Negative siRNA – Qiagen) were transfected using Lullaby transfection reagent according to manufacturer’s instructions.

For CrispR /Cas9 mediated knock out, sgRNA were selected using the MIT CrispR designing tool (http://crispr.mit.edu/). Annealed oligonucleotides were cloned into pLentiCrispR (pXPR_001) as described by ^14^. Briefly, HEK293T cells were seeded at 1.5 x 10^6^ cells/Ø10cm dish and transfected the day after. 10 μg of pLentiCrispR (empty vector or containing one of the following CrispR targeting sequence CrispR#1: 5’-CACCGGGTGCAGTTGTTCCACTAGT -3’or CrispR#2: 5’-

CACCGCCTGCTCTCGCTCTAGATGC -3’, were mixed with 7.5 μg of pSPAX2 and 4 μg of pVSVG in a final volume of 440μL of sterile water, and complemented with 500μL 2X HBS and 120mM CaCl_2_. Solution was incubated 30min at 37°C before being added onto HEK293T cells. Medium was removed 24h after transfection and replaced by 6mL of 20% FBS DMEM. Meanwhile, recipient cells were plated at 1 x 10^6^ cells/Ø10cm dish. The day after, supernatants were filtered through a

0.45 μm pore membrane and mixed with 25 μg of hexadimethrine bromide (4.2

μg/mL final) before infecting recipient cells. Infection was repeated the next day and stably transfected cells were selected using 1μg/mL of puromycin.

### FRET imaging of Mammalian Cells

The Rac1-Raichu-mTq2-sREACH probe will be described in ^15^. Cells were transfected with the probe, plated the day after on laminin and imaged. FRET images were acquired with the Nikon FLIM/TIRFsystem Z6014 microscope equipped with a Plan Apochromat 63x/1.45 oil objective and a 465nm LED. Dishes were fit in a 37°C heated chamber perfused with 5% CO_2_. FRET efficiency was calculated by standardizing the probe lifetime to the average lifetime of the donor alone as follows:

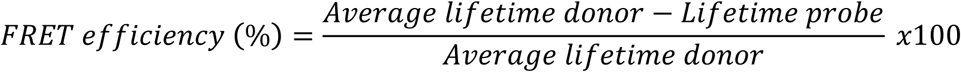

### Immunoprecipitation and Western Blotting of Mammalian Cells

Lysates were collected on ice by scraping cells in Ripa Buffer (150mM NaCl, 10mM Tris-HCl pH7.5, 1mM EDTA, 1% Triton X100, 0.1% SDS, 1X protease and phosphatase inhibitors) and centrifuged 10 min at 15000 rpm and 4°C. Protein concentration was measured at OD_600nm_ using Precision Red.

20-40μg of protein were resolved on a NuPAGE Novex 4-12% Bis Tris gels and transferred onto a nitrocellulose membrane using the BioRad system.

Membranes were blocked in 5% non-fat milk in TBS-T (10 mM Tris pH 8.0, 150 mM NaCl, 0.5% Tween 20) during 30min before overnight incubation with primary antibodies at 4°C. Membranes were washed 3x5 min in TBS-T and incubated 1h with Alexa-Fluor conjugated secondary antibodies. Blots were then washed 3x5 min and revealed using the LiCor Odyssey CLx.

All images were then analysed using Image StudioLite v.5.2.5.

### Immunofluorescence of Mammalian Cells

Cells were collected and plated onto sterile Ø13mm glass coverslips. Coverslips can be coated overnight at 4°C with 10μg/mL of rat tail collagen I, 10μg/mL fibronectin or 10μg/mL laminin diluted in PBS. Coating was washed out 3 times in PBS before seeding cells. Cells were allowed to adhere overnight (uncoated) or 4h (coated coverslip) and fixed with 4% paraformaldehyde for 10 min, permeabilised (20mM Glycine, 0.05% Triton X100) for 10 min and non specific binding was reduced by incubating coverslips 30 min in 5% BSA-PBS. Primary and secondary antibodies were diluted in blocking buffer and incubated 1h in a dark and humidified chamber. Coverslips were washed twice in PBS and once in water before being mounted on glass slides using ProLong Gold antifade reagent. Images were taken using an inverted Olympus FV1000 confocal microscope using a Plan Apochromat N 63x/1.40 oil SC or a Uplan FL N 40x/1.30 oil objective.

Images were processed and analysed using Fiji software (ImageJ v1.48t) ^16^.

### Random Migration Assay for Mammalian Cells

6-well glass bottom plates were coated overnight at 4 degrees with 10μg/mL of collagen I or fibronectin diluted in PBS. Coated dishes were washed in PBS. 1x10^5^ cells were plated and imaged every 10 min for 17 h using a Nikon TE2000 microscope, PlanFluor 10x/0.30 objective and equipped with a heated CO_2_chamber. Images were analysed using Fiji software^16^ (ImageJ v1.48t). Individual cells were tracked using the mTrackJ plugin, and spider plots were generated using the chemotaxis and migration tool plugin (v. 1.01).

### Inverted Invasion Assay of Mammalian Cells

We used a modified inverted invasion assay ^4, 17^ with Matrigel diluted 1:1 in ice cold PBS and 100 μL allowed to polymerize 30min at 37°C in each transwell.

Cells were diluted at 4x10^5^ cells/mL. 100 μL were plated as a confluent layer onto the underside of the filter. Transwells were then carefully covered with the base of the 24-well plate such that it contacts the droplet of cell suspension and left 2h in the incubator, allowing cell attachment. Transwells were washed into 3 x 1mL of serum free medium. The upper chamber contained 10% FBS supplemented medium, whereas the bottom chamber contained serum free medium. Cells migrated for 3 ½ days and stained 1h at 37 °C with 4 μM of Calcein AM (Life Technologies). Serial optical sections of 15μm interval were obtained with an Olympus FV1000 confocal microscope, UplanSApo 20x/0.74 objective.

Images were analysed using Fiji software^16^ (ImageJ v1.48t)

### *Dictyostelium discoideum* Cells

Axenic *D. discoideum* strains Ax3 was used as wildtype. *CYRI* knockout cells were generated in Ax3 genetic backgrounds. Ax3-derived *napA* KO cells are described elsewhere ^18^. Cells were grown in HL5 medium (Formedium) with 100U/ml penicillin and 100 μg/ml streptomycin in Ø10cm plastic Petri dishes and incubated at 21°C.

### *Dictyostelium discoideum* GFP-Trap with Formaldehyde Crosslinking

Cells were collected in PBS and lysed by adding ice-cold 3x lysis/crosslinking buffer (1x buffer: 20 mM HEPES pH 7.4, 2 mM MgCl_2_, 3% formaldehyde, 0.2% Triton X-100). After 5 min with gentle agitation at 4 °C, formaldehyde was quenched for 10min on ice using 1.75M Tris pH8.0. Samples were centrifuged at 22,000g for 4 min at 4 °C. Pellet was successively washed and resuspended with 1mL of ice cold quenching buffer (0.4 M Tris pH 8.0, 0.2% Triton X-100), wash buffer A (100 mM HEPES pH 7.4, 2 mM MgCl_2_, 0.2% Triton X-100) and wash buffer B (100 mM HEPES pH 7.4, 2 mM MgCl_2_), with 3 min centrifugation step between washes. Final resuspension was performed using 1mL of ice-cold RIPA buffer (50 mM Tris HCl pH 8.0, 150 mM NaCl, 0.5% Triton X-100, 0.5% sodium deoxycholate, 0.15% SDS, 5 mM EDTA, 2mM DTT) and incubated 1h at 4 °C with gentle agitation. Supernatants were mixed with pre-equilabrated GFP-Trap beads (Chromotek) following manufacturer’s protocole. Beads were washed 3 times with 50 mM Tris HCl pH 8.0, 150 mM NaCl, 5 mM EDTA followed by 1 wash with 10 mM Tris. Samples were eluted after incubation with 2x SDS loading buffer and heating 10min at 70°C before loading on a SDS-PAGE.

### *Dictyostelium discoideum* GFP-NAP1 ‘in gel’ Proteolytic Digestion – Mass Spectrometry Analysis

Eluates from GFP-NAP1 immunoprecipitation were separated by SDS-PAGE and stained with Coomassie blue. Each gel lane was divided in 6 slices and digested^19^. Tryptic peptides from in gel digestions were separated by nanoscale C_18_ reverse-phase liquid chromatography using an EASY-nLC II (Thermo Fisher Scientific) coupled online to a Linear Trap Quadrupole - Orbitrap Velos mass spectrometer (Thermo Scientific) and desalted using a pre-column C18 NS-MP-10 100μm i.d. x 0.2cm of length (NanoSeparations). Elution was at a flow of 300 nl/min over a 90 min gradient, into an analytical column C18 NS-AC-11 75μm i.d. x 15 cm of length (NanoSeparations). For the full scan a resolution of 30,000 at 400 Th was used. The top ten most intense ions were selected for fragmentation in the linear ion trap using Collision Induced Dissociation (CID) using a maximum injection time of 25 ms or a target value of 5000 ions. MS data were acquired using the XCalibur software (Thermo Fisher Scientific).

Raw data obtained were processed with MaxQuant version 1.5.5.1 ^20^ and Andromeda peak list files (.apl) generated were converted to Mascot generic files (.mgf) using APL to MGF Converter [http://www.wehi.edu.au/people/andrew-webb/1298/apl-mgf-converter.]. Generated MGF files were searched using Mascot (Matrix Science, version 2.4.1), querying dictyBase ^21, 22^(12,764 entries) plus an in-house database containing common proteomic contaminants and the sequence of GFP-NAP1. The common contaminant and reverse hits (as defined in MaxQuant output) were removed.

Mascot was searched assuming trypsin digestion allowing for two miscleavages with a fragment ion mass tolerance of 0.6 Da and a parent ion tolerance of 15 ppm. The iodoacetamide derivative of cysteine was specified in Mascot as a fixed modification, and oxidation of methionine and phosphorylation of serine, threonine and tyrosine were specified in Mascot as variable modifications. Scaffold (version 4.3.2, Proteome Software) was used to validate MS/MS based peptide and protein identifications. Peptide identifications were accepted if they could be established at greater than 95.0% probability as specified by the Peptide Prophet algorithm, resulting in a peptide false discovery rate (FDR) of 0.63% ^23^. For label-free quantification, proteins were quantified according to the label-free quantification algorithm available in MaxQuant ^23^.

Significantly enriched proteins were selected using a Welch-test analysis with a 5% FDR.

### Generation and Validation of *CYRI* knockout and Overexpressing *Dictyostelium discoideum*

Standard methods were used for construction of all Dictyostelium knockout and re-expression vectors^21, 22^. A linear CYRI knockout construct (2758 bp in length), which consisted of a blasticidin resistance (Bsr) cassette flanked by sequences matching 5’ and 3’ regions in the CYRI (DDB_G0272190 identifier at dictybase.org) gene locus (18pb cross-over), was made by PCR amplification using the following primers 5’-GTGACATTAAGGCAATAATAATAATAATAGAATTC-3’ and 5’-GTCTAGGAGCTCAGACGCCATCATAAGCAGTTCTAC-3’ (5’ arm), and 5’-GTGACAG TGCGTACTGGCCTCATATTATCGTCGTAC-3’ and 5’-TGGTGGATGATGGTATTTGGTGATTGTGATGTTTTC-3’ (3’ arm). PCR-amplified arms were combined with the Bsr cassette in a final PCR reaction using the following primers: 5’-GTGACATTAAGGCAATAATAATAATAATAGAATTC-3’ and 5’-TGGT

GGATGATGGTATTTGGTGATTGTGATGTTTTC-3’ and DNAs were purified using Phenol chloroform and ethanol precipitation. Knockout clones were screened/validated by PCR, with primers-5’-GAGGTCCAGC TATCATCGA-3’ and 5’-TCACATGTATCTTATGATAATG-3’. *CYRI* knockout yield a 2450 bp PCR product, random integrants (clones with a KO construct integration elsewhere in the genome) and wild-type yield a 1983 bp PCR product.

Vectors for expression of untagged CYRI was obtained by sub-cloning CYRI’s geomic coding region into pDM358 ^24^. In this construct, CYRI’s open reading frame (ORF) is placed in an expression cassette driven by the strong *act15* promoter. According to dictyExpress, ^25^, endogenous act15 gene is more highly expressed than CYRI gene in vegetative Ax2 and Ax4 cells. Additionally, the CYRI expression vector is likely to be present in multiple extrachromosomal copies in any transformant cell ^24, 26^, driving high CYRI expression levels. A REMI (non extra-chromosomal) vector was derived from this by removal of the Dictyostelium plasmid propagation genes and re-ligation of the vector backbone. This construct, while still having a strong promoter, is expected to be present in just one copy per cell.

### Transformation of *Dictyostelium discoideum*

3.0 x 10^7^ cells/transformation were first centrifuged (3 min, 330 *g*, 4°C), washed with 10 ml ice-cold electroporation buffer (E-buffer; 10 mM sodium phosphate buffer pH 6.1, 50 mM sucrose), and resuspended in 400 μl ice-cold E-buffer. Cells were transferred into an ice-cold 0.2 cm electroporation cuvette and incubated 5 min with 0.5-1.0 μg of DNA on ice. Cells were electroporated (BTX-Harvard Apparatus ECM 399) at 500V, giving a time constant of 3-4ms. Cells were immediately transferred to HL5 medium in Petri dishes. Appropriate selection (50 μg/ml hygromycin or 10 μg/ml G418) was added the next day. For REMI transfections, 10 μg of linearized DNA and 50 U of restriction enzyme were used, in 0.4cm cuvettes with a Bio-Rad Gene Pulser II set at 1.2kV and 3μF.

### *Dictyostelium discoideum* Under-agarose Chemotaxis Assay

This assay is based on a previous study ^27^. Surface of the Ø35mm glass bottom dish (MatTek) was coated with 10 mg/ml BSA for 10 min, washed with dH_2_O and dried for 5 min inside a laminar flow cabinet. 0.4% w/v SeaKem GTG agarose in SIH medium (Formedium) containing 10 μM folate was poured and set for 1h. A well was cut in the agarose and 2x10^6^ cells placed in it. After 3-4h cells had migrated sufficiently into the agarose and were imaged by Phase contrast and DIC microscopy was performed with a Nikon Eclipse TE2000-E microscope system equipped with a QImaging RETIGA EXi FAST 1394 CCD camera and a pE-100 LED illumination system (CoolLED) at 525 nm. A 20×/ 0.45 NA Ph1 objective and a 60×/1.40 NA apochromatic DIC objective were used for phase contrast and DIC, respectively. Imaging was controlled through the μManager 1.4.9 software. All microscopy was carried out at room temperature and images were analysed with ImageJ/Fiji 1.49i.

Pseudopod rate and split frequency was analysed from the DIC movies and manually quantified frame by frame. For analysis of cell Circularity, speed and migration parameters automated tracking plugins were developed for ImageJ. More information will be supplied upon reasonable request, but the code used was as follows:

~~~
run(“Convert to Mask”, “method=IsoData background=Dark calculate black“);
run(“Invert”, “stack“);
run(“Analyze Particles…”, “size=400-Infinity pixel circularity=0.00-1.00
show=Masks in_situ stack“);
run(“Fill Holes”, “stack“);
run(“Set Measurements…”, “area centroid shape stack redirect=None
decimal=3“);
run(“Analyze Particles…”, “size=0-Infinity circularity=0.00-1.00 show=Nothing
display clear summarize stack“);
~~~

### Statistical Analysis

Data sets were analysed using Prism5 v5.0c and tested for normality. Differences between groups were then analysed using the appropriate statistical test, mentioned in each figure legend. Error bars represent standard error of the mean (SEM). Significance levels are given as follow: ns: p >0.05; *: p ≤0.05; **p≤0.01; ***: p ≤0.001. Cochran-Mantel-Haenszel test and Likelihood ratio test with Bonferroni correction test were generated using R software and *p*-value are mentioned when appropriate.

